# cGAS-STING drives alveolar epithelial cell dysfunction in cigarette smoke-induced lung injury

**DOI:** 10.64898/2026.06.16.732582

**Authors:** Rocío Fuentes-Mateos, Pratama A. Saputra, I. Sophie T. Bos, Vicky Verschut, Iris C. Gorter, Justina C. Wolters, Barbro N. Melgert, Reinoud Gosens

## Abstract

Chronic inflammation induced by cigarette smoke (CS) plays a central role in the pathogenesis of chronic obstructive pulmonary disease (COPD), but its impact on lung epithelial progenitor function and regenerative capacity remains incompletely understood. Here, we combined *in vivo* and *in vitro* approaches to dissect how CS exposure and subsequent inflammatory insults shape epithelial repair dynamics. A 6-week whole-body CS exposure model in mice induced lung function impairment and altered gene expression profiles in alveolar epithelial cells, prominently activating interferon (IFN)-related pathways and the cGAS-STING axis. Alveolar epithelial cells from CS-exposed mice generated a similar number, but larger organoids with reduced alveolar differentiation compared to air-exposed mice. Notably, these cells obtained from CS-exposed mice displayed resistance to IFNγ-induced suppression of organoid growth, contrasting with the strong inhibitory effect of IFNγ observed in controls. This phenotype was recapitulated in a two-hit *in vitro* model using cigarette smoke extract (CSE), in which chronic CSE exposure impaired regeneration and differentiation while inducing resistance to IFNγ. Gene expression and proteomic analyses revealed upregulation of Zbp1, Irf7, and other upstream IFN regulators, correlating negatively with alveolar differentiation potential. Inhibition of the cGAS-STING pathway with RU.521 partially rescued organoid formation, increased proliferation, and alveolar differentiation. Together, our data reveal that CS exposure alters the alveolar epithelial landscape, inducing a stress-adapted, IFNγ-resistant state that compromises alveolar regeneration, with cGAS-STING activation as a key driver of early CS-associated alveolar type 2 cell dysfunction. These findings provide new insight into how chronic inflammation reshapes progenitor cell function during early lung injury.

## Introduction

Chronic obstructive pulmonary disease (COPD) is a prevalent and progressive respiratory condition characterized by heterogeneous expression of persistent bronchitis and structural remodeling, as well as impaired tissue repair associated with emphysema, ultimately leading to a decline in lung function ^1,2^. Cigarette smoke (CS) is the primary risk factor, largely due to its high oxidant effect and the persistent lung and systemic inflammation it triggers. CS-derived oxidants directly damage lung cells and tissues, deplete antioxidant defenses, and promote chronic inflammation. Notably, this inflammatory response can persist long after smoking cessation, sustained by altered immune responses, retained particulates in the lung, and epigenetic modifications driven by oxidative stress ^3–5^.

In addition to causing direct tissue damage, CS-derived oxidants and inflammatory mediators contribute to the destruction of lung parenchyma and the senescence of alveolar type II (AT2) and mesenchymal stem cells. This promotes the loss of alveolar epithelial integrity and the development of emphysema ^1,5–7^. The continued secretion of inflammatory mediators, coupled with an ineffective repair response, accelerates lung function decline in patients with COPD.

While oxidative stress and inflammation are well-established contributors to COPD pathogenesis, the molecular mechanisms through which CS impairs cellular regeneration remain incompletely understood. AT2 cells, which are central to alveolar repair, are particularly sensitive to external stimuli from surrounding cells, alterations in the extracellular matrix, and interactions with supportive mesenchymal cells ^8–10^.

Although acute inflammation can promote tissue repair, chronic or unresolved inflammation has the opposite effect, impairing progenitor cell function and contributing to sustained tissue damage. Acute inflammatory responses are typically triggered by injury and activate pathways that support healing and restoration of homeostasis. Importantly, this process is tightly regulated and includes the production of immunosuppressive cytokines such as IL-10 and TGF-β, which help resolve inflammation and return the tissue to a homeostatic state ^11^.

In contrast, persistent inflammation represents a dysregulated response driven by continuous stimuli, such as aging, cigarette smoke exposure, metabolic disorders, or chronic antigenic stimulation ^12^. Under these conditions, immune cells remain activated at the site of injury, leading to excessive production of proteases, reactive oxygen species, and pro-inflammatory cytokines, and promoting a sustained shift toward a pro-inflammatory phenotype ^11^.

Several inflammatory cytokines, including TNFα, IL-1β, IL-6, and IL-8, play dual roles in lung repair, particularly during COPD exacerbations ^13^. These cytokines not only originate from inflammatory cells but are also secreted by senescent cells, linking chronic inflammation to accelerated lung aging. Interestingly, the presence of these cytokines in patients has been linked to exacerbation periods ^13^. However, the addition of the aforementioned cocktail of cytokines to an *in vitro* model of lung regeneration increased epithelial proliferation but altered differentiation ^14^. IL-1β treatment alone also showed a positive effect on epithelial cell proliferation in an organoid model, but a detrimental effect when the supportive fibroblasts were pre-treated ^15^. In addition to these cytokines, interferon-gamma (IFNγ) has emerged as a key regulator of tissue remodeling and emphysema progression in COPD. Tissue-resident lymphocytes are the primary source of IFNγ in COPD lung tissue ^16^. While it appears to be beneficial at low doses ^17^, proliferative AT2 cells appear particularly vulnerable to its effects ^16^. This highlights that not only the dose or duration of the inflammatory stimuli is key in the regenerative response, but also the recipient cell, and further elucidating how the inflammatory niche affects lung epithelial repair is crucial.

In this study, we investigated how chronic exposure to cigarette smoke changes the regenerative capacity of the AT2 cells and how pre-existing inflammation shapes the epithelial response to secondary inflammatory stimuli. Using a combined dataset with RNAseq and proteomics data of the effect of chronic *in vivo* cigarette smoke exposure in epithelial cells, along with *in vitro* organoid studies, we show that CS exposure alters the alveolar epithelial landscape, inducing a stress-adapted, IFNγ-resistant state that compromises alveolar regeneration, with cGAS-STING activation as a key driver of early CS-associated alveolar type 2 cell dysfunction.

## Materials and Methods

Detailed methods are provided in the Supplementary Material.

### Animal experiments and cigarette smoke exposure

All animal experiments were approved by the institutional animal care committee of the University of Groningen in accordance with national guidelines (license AVD10500209120). Male C57BL/6 mice (8–12 weeks) were randomly assigned to air (AIR) or cigarette smoke (CS) (n = 8/group), and exposed whole-body to the smoke of 3R4F research cigarettes, as previously described in ^18^ for 6 weeks to induce chronic low-grade lung inflammation, while control animals were exposed to filtered air. Lung function was assessed 16 hours after the final exposure.

### Lung function measurements

Pulmonary function was measured using a FlexiVent system following tracheostomy and mechanical ventilation as described in ^19^. Parameters, including elastance, compliance, and forced expiratory volume, were obtained using standardized perturbation maneuvers.

### Isolation of epithelial progenitor cells and macrophages

Lung epithelial cells were isolated by enzymatic digestion followed by magnetic sorting to obtain CD45⁻/CD31⁻/CD326⁺ (Epcam+) epithelial progenitor cells as described in ^19^, a population enriched for AT2 cells ^14,20^.

Macrophages were isolated from lung tissue using the CD31+/CD45+ fraction, as described previously ^21^.

### Organoid culture and in vitro cigarette smoke exposure

Epcam⁺ cells were co-cultured with mitotically inactivated CCL-206 murine lung fibroblasts in a 3D Matrigel-based organoid system as described in ^19^. Organoids were treated with 5 % cigarette smoke extract (CSE) for 14 days ^18,19^. A two-hit model was established by dissociating organoids and re-plating cells in the presence of secondary inflammatory stimuli, including Poly(I:C), LPS, cytokine mixtures, or IFNγ (**Table 1**). Organoid number, size, and differentiation were quantified after 14 days.

**Table 1:**
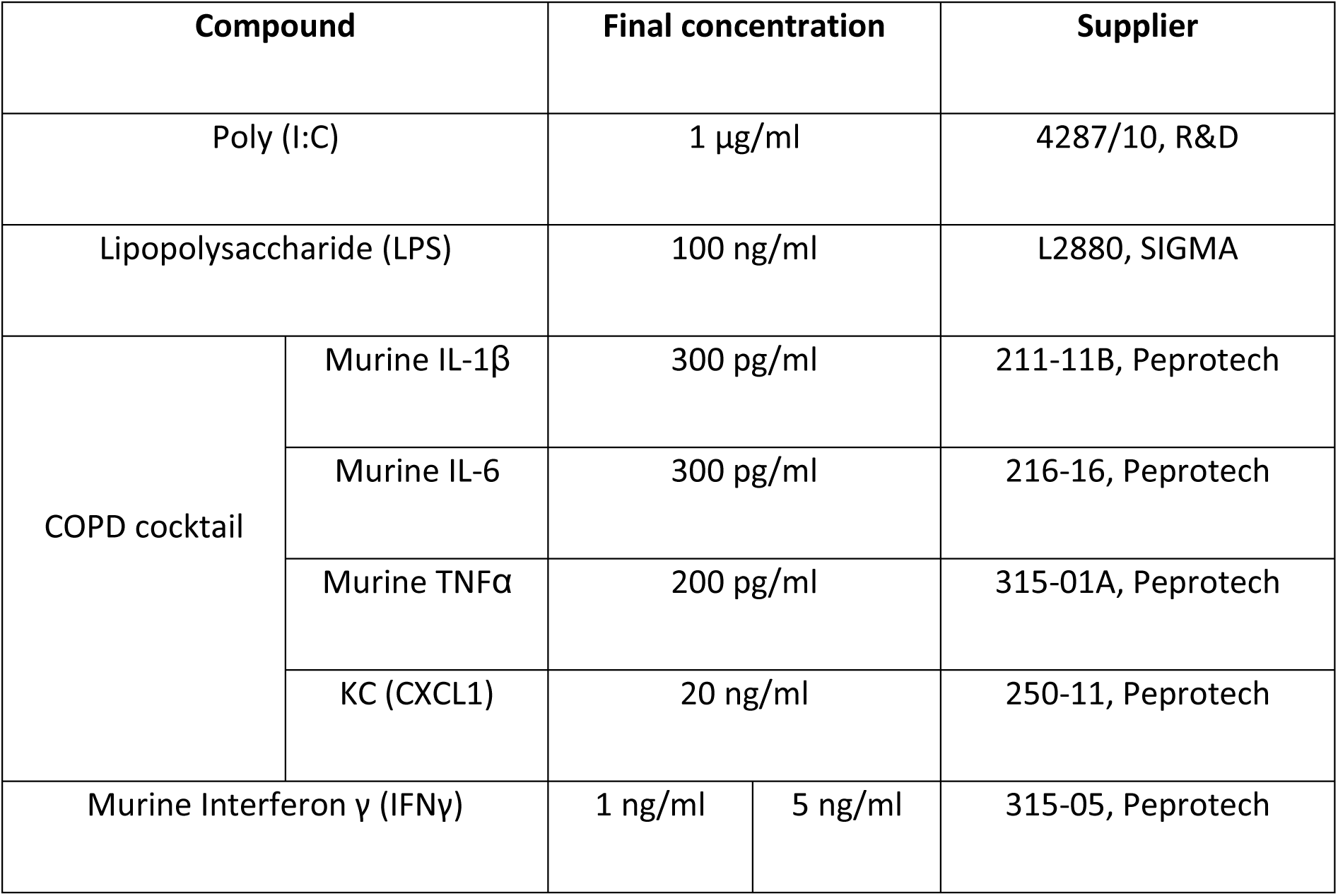
Overview of the compounds employed in the organoid assays, including supplier information and catalog numbers.

### Organoid dissociation and separation of epithelial and mesenchymal fractions

For organoid resorting, Epcam+ cells and CCL-206 fibroblasts were seeded as described in ^19^. Organoids were treated with 5 % CSE and 3 µM of RU.521, or their respective vehicles for 14 days, dissociated and separated into epithelial (CD326⁺) and mesenchymal (CD326⁻) fractions using magnetic-activated cell sorting for downstream RNA-seq analysis.

### RNA sequencing and pathway analysis

RNA sequencing was performed on freshly isolated epithelial cells (CD326+) and macrophages, as well as CD326^+^ and CD326^-^ (CCL-206) fractions from resorted organoids. Differential expression analysis was conducted using DESeq2. Pathway enrichment was assessed using gene set enrichment analysis (GSEA) and gene set variation analysis (GSVA). Correlation analyses were performed using Pearson’s correlation.

### Proteomics

Label-free quantitative proteomics was performed on freshly isolated epithelial cells, and differentially regulated proteins were identified and integrated with transcriptomic data as previously described in ^22^.

### Statistical analysis

Data are presented as mean±SEM or median with interquartile range, as appropriate. Normality was assessed using multiple tests, and parametric or non-parametric tests were applied accordingly. Comparisons between two groups were performed using unpaired or paired two-tailed Student’s t-tests or Mann–Whitney/Wilcoxon tests. For multiple group comparisons, one-way or two-way ANOVA was used. Sample sizes, biological replicates, and statistical tests are indicated in the figure legends. Statistical significance was defined as p < 0.05. Analyses were performed using GraphPad Prism (version 10) and R (version 4.2.0).

## Results

### Cigarette smoke exposure for 6 weeks alters lung function

To investigate the impact of chronic low-grade inflammation induced by cigarette smoke (CS), we employed a 6-week whole-body CS exposure mouse model (Fig. 1A). During the exposure period, control mice showed a steady increase in body weight, whereas CS-exposed mice failed to gain weight (Fig. 1B). Lung function analysis revealed a significant upward and leftward shift in the pressure–volume curve in CS-treated animals (Fig. 1C), accompanied by decreased lung elastance and increased compliance (Fig. 1D). Moreover, CS exposure led to a significant increase in forced expiratory volume at 0.1 seconds (FEV₀.₁), but no significant changes in the ratio FEV₀.₁/FVC or inspiratory capacity (Fig. 1D). Hematoxylin and eosin (H&E) staining showed a trend toward alveolar enlargement, although no significant difference in mean linear intercept (LMI) was detected (Fig. 1E). Together, these findings demonstrate that six weeks of CS exposure are sufficient to impair pulmonary function without overt structural alterations, reflecting an early disease stage in which emphysema is not yet established.

**Figure 1:**
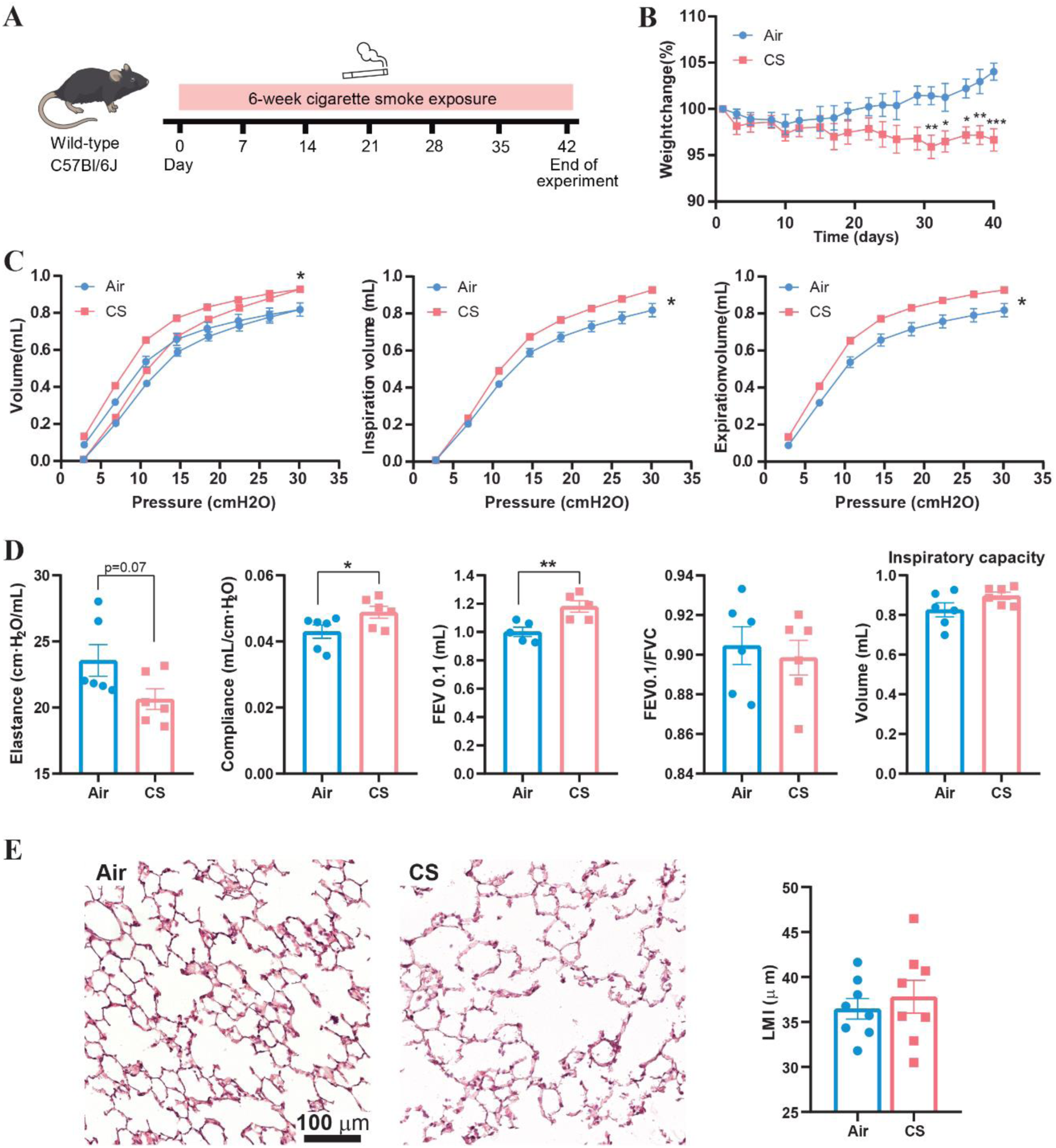
Six-week cigarette smoke exposure induces early lung function impairment and structural changes. (A) Experimental design showing the *in vivo* 6-week cigarette smoke (CS) exposure murine model. (B) Graph depicting the weight changes over the 6-week exposure period, expressed as % of initial weight. Data represent mean ± SEM of 8 mice per group. Statistical analysis was performed using two-way ANOVA followed by Sidak’s multiple comparisons test, * p<0.05, ** p< 0.01, *** p<0.001. (C) Flexivent pressure-volume (PV) curves for inspiration and expiration, including separate inspiration and expiration curves. The data shown are the means for 6 mice per group. Pressure-volume curves were analyzed using nonlinear regression (Gaussian model). Differences between datasets were assessed using an extra sum-of-squares F test, * p<0.05. (D) Flexivent lung function parameters: tissue elastance and compliance; forced expiratory volume at 0.1 seconds (FEV₀.₁); FEV₀.₁/forced vital capacity (FVC) ratio; and inspiratory capacity. Data represent mean ± SEM of n=6 independent mice per group. Statistical analysis was performed using a paired two-tailed Student’s t-test, * p<0.05, ** p< 0.01. (E) Representative Hematoxylin-Eosin (H&E) staining of lung sections from Air or CS mice (left) and corresponding linear mean intercept measurements (LMI, right). Scale bar 100 µm. Data represent mean ± SEM of 8 mice per group. Statistical analysis was performed using a paired two-tailed Student’s t-test.

### cGAS-STING pathway and interferon response activation upon in vivo CS exposure

To explore the impact of CS exposure on epithelial progenitor cells (Epcam⁺), we isolated CD45⁻/CD31⁻/Epcam⁺ cells from mouse lungs and performed bulk RNA-sequencing and proteomic analyses. Of these approaches, transcriptomics had the largest feature depth and was therefore used for our initial analyses. Since the Epcam⁺ population is not composed exclusively of AT2 cells, we performed cell-type deconvolution using GSE124872^23^ as a reference and adjusted our RNA-seq data according to the estimated AT2 proportion (Suppl. Fig. 1A). Deconvolution analysis revealed no significant reduction in AT2 cells, although a decreasing trend was observed. In contrast, AT1 cells appeared underrepresented, while goblet cell signatures were increased (Suppl. Fig. 1A). Correction for the AT2 proportion had minimal impact on global gene expression patterns, as shown by the MA plot (Suppl. Fig. 1B), consistent with the enrichment of AT2 cells within the Epcam⁺ population. Principal component analysis (PCA) revealed that epithelial cells from CS-exposed mice clustered separately from controls (Fig. 2A), indicating distinct transcriptional profiles. Differential expression analysis identified 1.786 differentially expressed genes (DEGs; padj < 0.05 and |log2 fold change| ≥ 0.58, corresponding to a ≥1.5-fold change), of which 1.227 were upregulated, and 559 were downregulated. Many of the highly upregulated genes were associated with inflammatory pathways (Fig. 2B).

**Figure 2:**
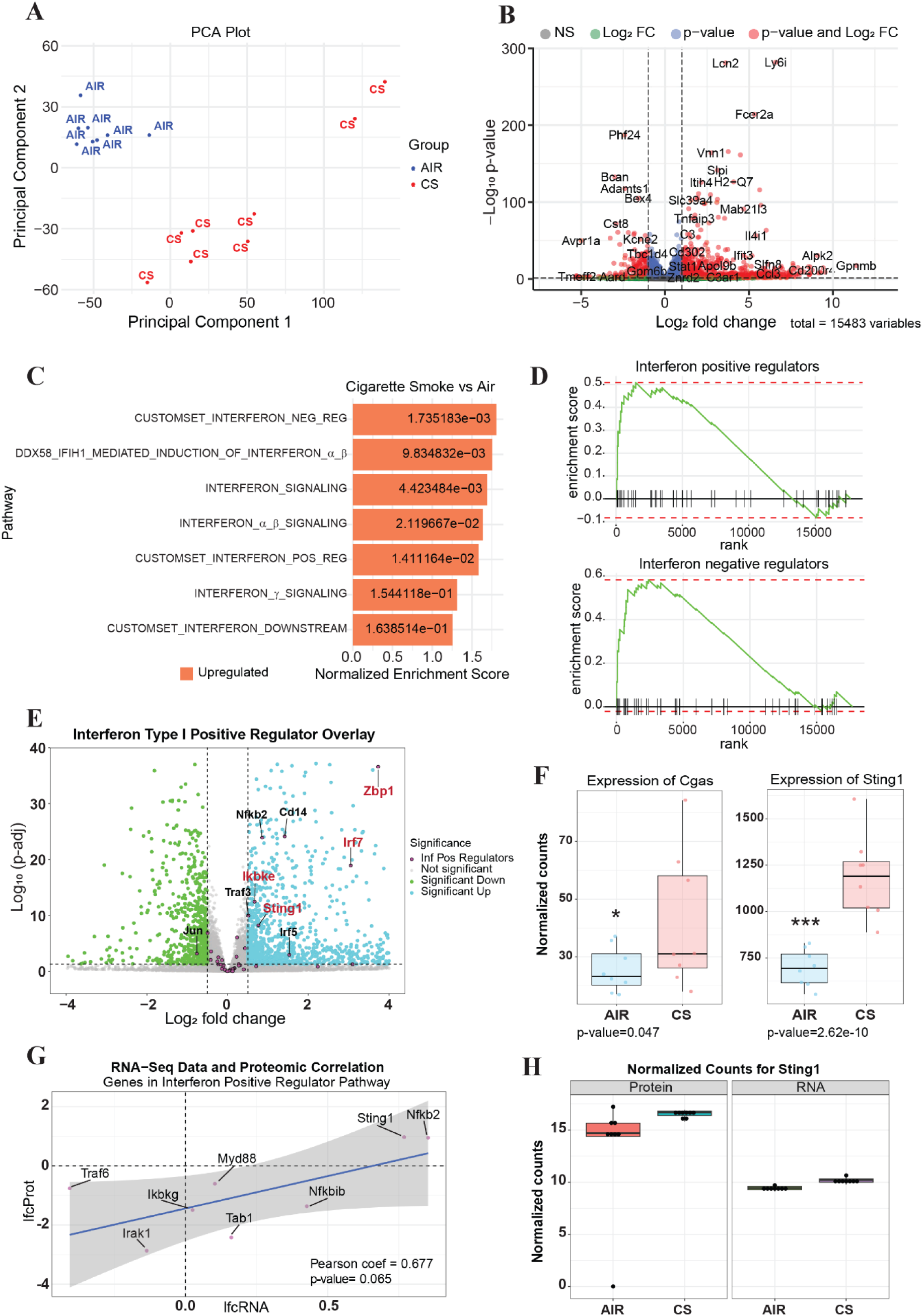
Cigarette smoke exposure activates interferon signaling via the cGAS-STING pathway in lung epithelial progenitors. (A) Principal component analysis (PCA) of CD45⁻/CD31⁻/Epcam⁺ cells from Air or CS-exposed mice, showing distinct clustering. (B) Volcano plot of differentially expressed genes (DEGs; padj < 0.05) of a total of 15.483 genes in Epcam⁺ cells. DEGS in CS *vs* Air samples are highlighted in red. (C) Gene set enrichment analysis (GSEA) of Reactome pathways showing enrichment of interferon signaling-related pathways in CS *vs* Air Epcam+ cells. p-values are indicated in the graph. (D) Representative GSEA plots for *Interferon positive regulators* and *Interferon negative regulators* pathways. (E) Volcano plot showing upregulation of cGAS-STING pathway components highlighted in red (Zbp1, Tmem173/Sting1, Ikbke, Irf7) in CS *vs* Air Epcam+ cells. (F) Normalized expression counts for *Mb21d1* (cGAS) and *Tmem173* (STING) in Air and CS samples. Data is represented as median with interquartile range and minimum-to-maximum values of n = 8 mice per group. Statistical significance was determined by differential expression analysis (DESeq2), and p-values are indicated in the graph. * p<0.05, *** p<0.001. (G) Proteomics validation showing the Pearson correlation between interferon-related gene expression (RNA-seq) and protein abundance (proteomics). Each point represents a gene, with positive regulators highlighted. Pearson correlation coefficient (r) is indicated. (H) Comparison of *Tmem173*/*Sting1* expression at the RNA (normalized counts) and protein (relative abundance) levels across experimental conditions.

Gene set enrichment analysis (GSEA) using the Reactome database demonstrated significant enrichment in pathways related to inflammation, viral responses, oxidative stress, and interferon signaling. Notably, in addition to “*Interferon Signaling,”* the pathways *“Interferon Positive Regulators*,” “*Interferon Negative Regulators*,” and “*DDX58/IFIH1-Mediated Induction of Interferon Alpha/Beta*” were also enriched in lung tissue of CS exposed mice (Fig. 2C), with GSEA plots showing upregulation of most genes within these interferon-related pathways (Fig. 2D). Proteomics analysis further supported these findings: of the 380 interferon-related genes identified by RNA-seq, 58 were also significantly changed in the proteomics dataset (Suppl. Fig. 2), showing strong concordance, particularly among positive regulators (Fig. 2G, H).

Among the top upregulated genes in CS-exposed mice was Z-DNA binding protein 1 (*Zbp1*) (Fig. 2E), a cytosolic sensor of nucleic acids known to regulate antiviral responses, inflammation, and cell death. Zbp1 is associated with nucleic acid-sensing pathways that converge on interferon signaling. Additional components of the cGAS-STING pathway, such as *Tmem173* (*Sting1*), *Ikbke* (*IKKε*), and *Irf7*, were also upregulated (Fig. 2E). In line with these findings, normalized expression counts for *Mb21d1* (cGAS) and *Tmem173* (STING1) were significantly increased in Epcam⁺ cells from CS-exposed lungs compared to controls (Fig. 2F). Notably, expression of interferon ligands was minimal or undetectable in Epcam⁺ cells (Suppl. Fig. 3A). In contrast, several interferon receptors, including *Ifnra2, Ifngr1, Ifngr2 and Il10rb* were significantly upregulated under CS conditions (Suppl. Fig. 3A). A similar pattern was observed in macrophages isolated from the same mice, which also showed low interferon gene expression but significantly higher expression of interferon receptors (*Ifnra1, Ifnra2, Ifngr2* and *Il10rb*) (Suppl. Fig. 3B). The cGAS-STING axis senses cytosolic DNA, not only of viral or bacterial origin but also derived from nuclear or mitochondrial damage. Together, these findings suggest that CS exposure is associated with activation of interferon-responsive signaling programs and increased interferon sensitivity, rather than robust endogenous interferon production, potentially linked to cGAS-STING activation secondary to DNA damage.

Consistent with the trends observed in the deconvolution analysis (Suppl. Fig. 1A), key AT2 markers, including *Abca3, Lamp3, Lpcat1, Sftpb,* and *Sftpc*, were reduced in CS samples compared to AIR controls at both the transcriptomic and proteomic levels (Suppl. Fig. 4), indicating a shift in epithelial cell composition and/or state.

### Epcam+ cells exposed in vivo to CS exhibit a decrease in in vitro alveolar differentiation and reduced sensitivity to IFNy

To investigate how chronic *in vivo* CS exposure affects the regenerative capacity of Epcam⁺ epithelial cells, we employed an *in vitro* 3D lung organoid model. This system recapitulates key stages of epithelial repair, including survival post-injury, proliferation, and differentiation. To mimic the inflammatory microenvironment of COPD exacerbations requiring lung repair, a second inflammatory stimulus was introduced to the organoid cultures (**Table 1**, M&M).

Interestingly, the 6-week CS exposure did not significantly impact organoid-forming efficiency (Fig. 3A, B). However, organoids derived from CS-treated mice were larger in size (Fig. 3A, B) and exhibited a reduction in alveolar differentiation, as evidenced by fewer alveolar-like organoids (Fig. 3C) and a significantly decreased proportion of SftpC⁺ organoids, accompanied by an increase in double-negative (DN, SftpC^-^/Acetylated-Tubulin^-^) organoids (Fig. 3D).

**Figure 3:**
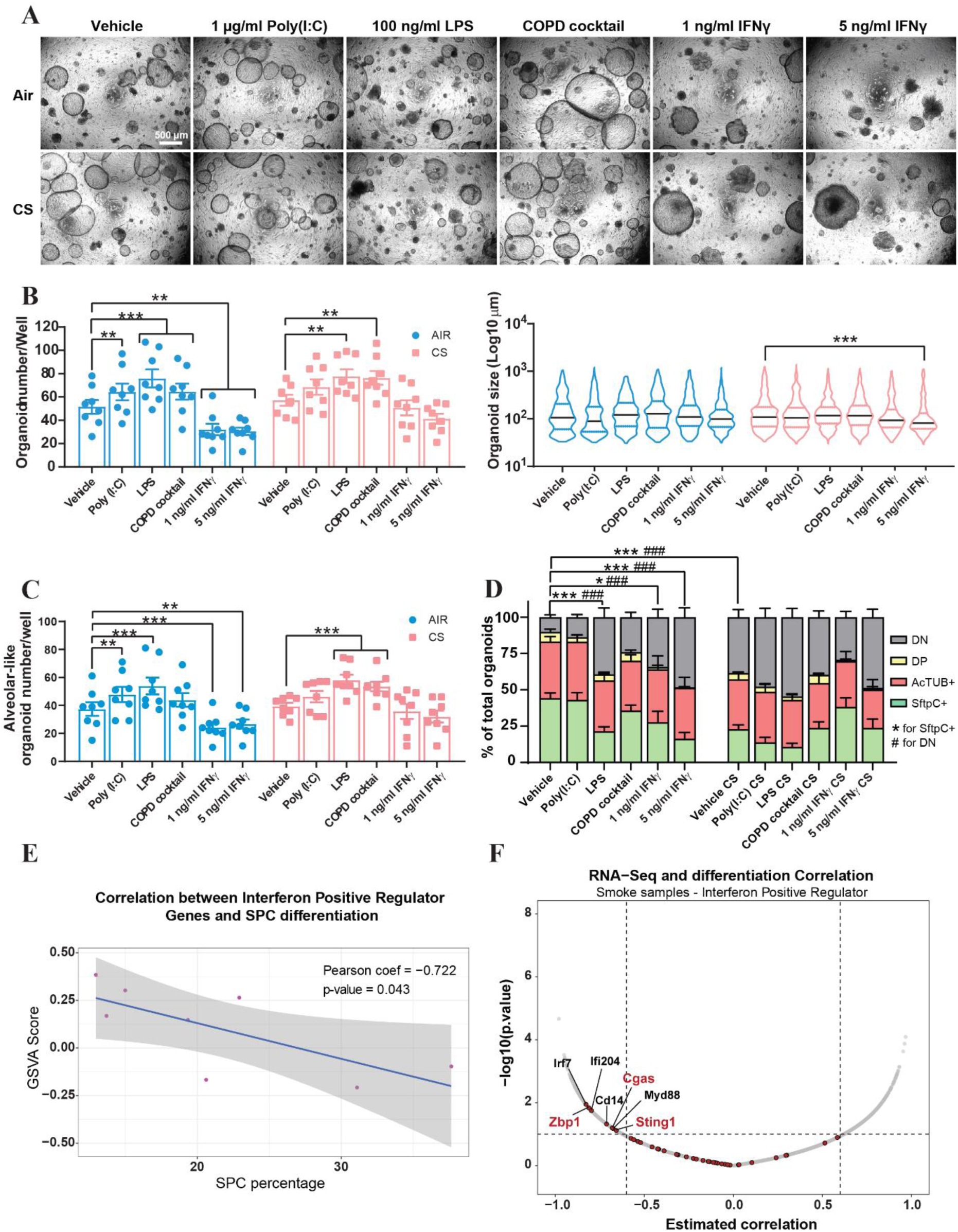
I*n vivo* cigarette smoke (CS) exposure impairs alveolar differentiation and alters epithelial progenitor response to secondary inflammatory stimuli in 3D lung organoids. (A) Representative images of organoids derived from *in vivo* Air or CS-exposed mice with or without the indicated secondary inflammatory stimuli. Scale bar 500 µm. (B) Quantification of organoid-forming efficiency (left) and organoid size (right) under the indicated conditions. For organoid number, data represent mean ± SEM of n = 8 mice, and statistical analysis was performed using two-way ANOVA followed by Sidak’s multiple comparisons test. For organoid size, data represent median ± quartiles of n = 8 mice, and statistical analysis was performed using Kruskal-Wallis followed by Dunńs multiple comparisons test. ** p< 0.01, *** p<0.001. (C) Quantification of total alveolar-like organoid number. Data represent mean ± SEM of n = 8 mice. Statistical analysis was performed using two-way ANOVA followed by Sidak’s multiple comparisons test, ** p< 0.01, *** p<0.001. (D) Quantification of percentage of Sftpc⁺ alveolar organoids, AcTub^+^ airway organoids, double-positive (DP, SftpC^+^/AcTub^+^) and double-negative (DN, SftpC^-^/AcTub^-^) organoids. Data represent mean ± SEM of n = 8 mice. Statistical analysis was performed using two-way ANOVA followed by Dunnett’s multiple comparisons test, * indicates differences in SftpC^+^, # indicates differences in DN. *p< 0.05, ***/### p< 0.001. (E) Pearson correlation between GSVA enrichment score for *IFN positive regulator* pathway and percentage of SftpC⁺ organoids per sample. Each point represents an individual biological sample, n=8 mice. The solid line indicates the linear regression fit. Pearson correlation coefficient (r) and p-value are shown. (F) Volcano plot showing gene-level correlations between *Interferon positive regulators* expression and organoid alveolar differentiation capacity, measured as the percentage of SftpC+ organoids. Each point represents a gene; the x-axis indicates the Pearson correlation coefficient, and the y-axis shows −log10(p-value). Genes with negative correlation coefficients are associated with reduced alveolar differentiation. Selected negatively correlated genes (*Tmem173*/*Sting1*, *Zbp1,* and *Mb21d1/cGas*) are highlighted in red.

Upon addition of secondary inflammatory stimuli, including Poly(I:C), LPS, and a COPD cytokine cocktail, organoid-forming efficiency increased in both AIR and CS groups (Fig. 3A, B). Nevertheless, SftpC⁺ organoids remained significantly reduced in the CS-derived samples (Fig. 3D), indicating persistent impairment in alveolar differentiation. Notably, IFNγ treatment severely disrupted organoid formation in AIR-derived cells but had less impact on the organoid-forming efficiency of CS-exposed cells (Fig. 3B). In contrast, organoid growth (size) was unaffected by IFNγ in AIR exposed cells, whereas IFNγ did cause a concentration-dependent reduction in organoid size in the CS-exposed cells (Fig. 3B). These findings suggest that prior *in vivo* CS exposure not only rewires the transcriptomic landscape, upregulating interferon-responsive genes, but also alters the functional response of epithelial progenitors to subsequent IFNγ exposure (Fig. 3C, D).

Given that Epcam⁺ cells used for RNA-seq and organoid experiments were derived from the same individual mice, we next assessed whether interferon signaling correlated with the *in vitro* alveolar differentiation potential. Using Gene Set Variation Analysis (GSVA), we computed enrichment scores for the “*IFN positive regulator*” pathway per sample and correlated these with the percentage of SftpC⁺ organoids. A significant (Pearson’s r=-0.722, p=0.043) negative correlation was observed: higher pathway enrichment was associated with lower alveolar differentiation capacity (Fig. 3E). This relationship was further supported by gene-level correlation analysis, visualized as a volcano plot (Fig. 3F), in which each point represents an individual gene within the *Interferon positive regulator* pathway. The x-axis shows the Pearson correlation coefficient between gene expression and the percentage of SftpC^+^ organoids, while the y-axis represents the statistical significance (-log10 p-value) of the correlation. Genes with more negative correlation coefficients are associated with lower alveolar differentiation. Among these, *Irf7* and *Zbp1* emerged as the most strongly negatively correlated genes, suggesting their potential role as suppressors of epithelial regeneration following CS exposure.

### In vitro cigarette smoke exposure distorts organoid growth, differentiation, and IFNγ response

Epcam⁺ cells isolated from mice exposed *to* CS *in vivo* are influenced not only by cigarette smoke itself but also by the complex lung inflammatory microenvironment and the actions of the immune system. To determine whether CS alone can directly impact epithelial cell behavior, we established a two-hit organoid model. In this system, cells were exposed to 5 % CSE *in vitro* for 14 days to mimic chronic exposure. After this period, organoids were dissociated into single cells to simulate injury and subsequently replated in the presence or absence of a secondary inflammatory stimulus (**Table 1**, Fig. 4A).

**Figure 4:**
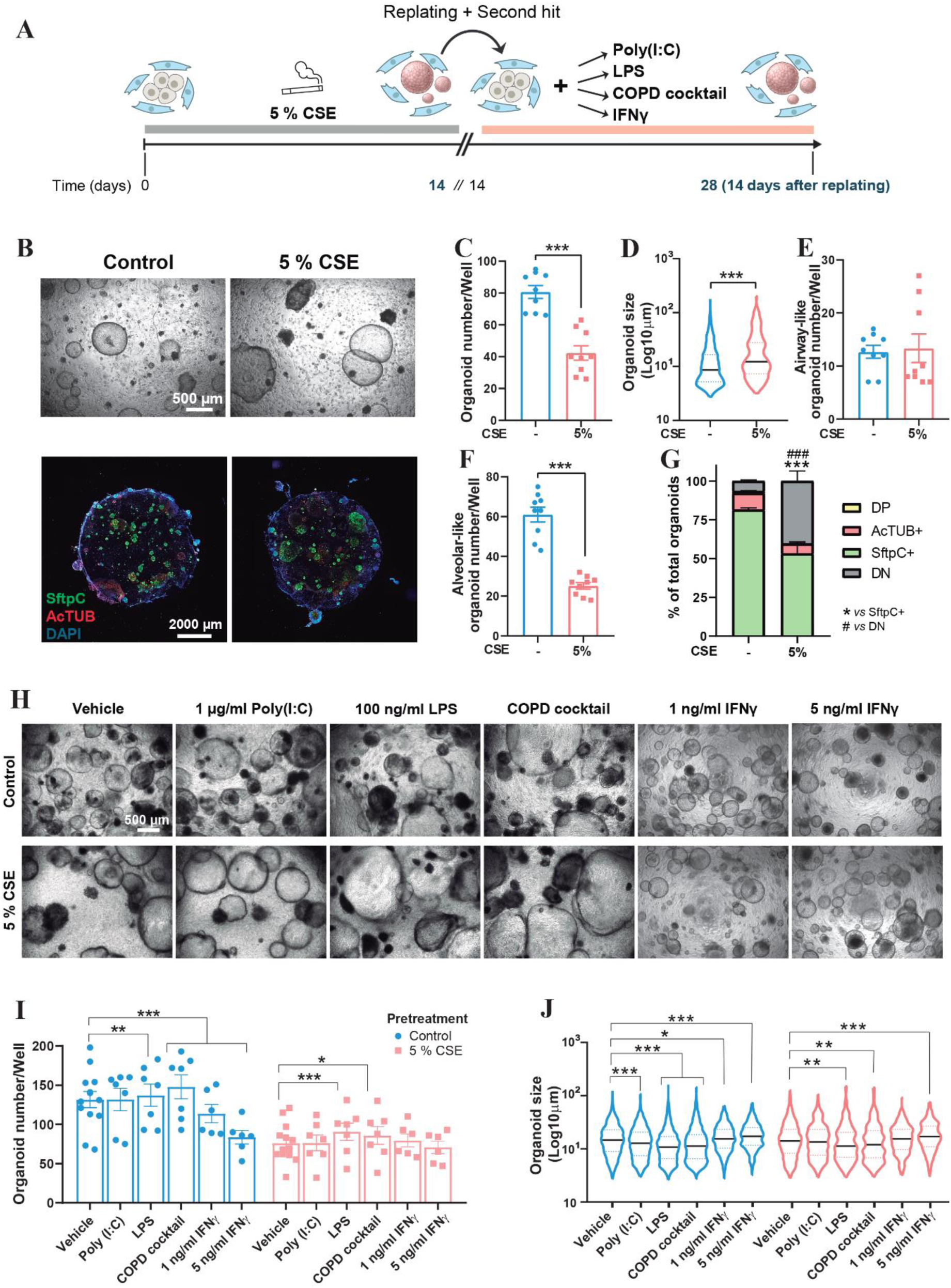
I*n vitro* cigarette smoke extract (CSE) exposure reduces alveolar regeneration, alters organoid growth, and modulates IFNγ sensitivity in a two-hit organoid model. (A) Schematic representation of the two-hit organoid model, including 14-day 5% CSE exposure followed by replating with or without secondary inflammatory stimulus. (B) Representative light microscopy images of organoids derived from control and *in vitro* 5 % CSE exposure before replating (Upper panels, scale bar 500 µm). Representative immunofluorescence images of organoids with SftpC^+^ (green) and AcTub^+^ (red) markers, counterstained with DAPI (blue) (Bottom panels, scale bar, 2000 µm). (C) Quantification of organoid-forming efficiency. Data represent mean ± SEM of n = 9 mice. Statistical analysis was performed using an unpaired two-tailed Student’s t-test, *** p<0.001. (D) Quantification of organoid size under the indicated conditions. Data represent median ± quartiles of n = 9 mice. Statistical analysis was performed using a Mann-Whitney two-tailed test, *** p<0.001. Quantification of total airway-like (E) and alveolar-like (F) organoids under the indicated conditions. Data represent mean ± SEM of n = 9 mice. Statistical analysis was performed using an unpaired two-tailed Student’s t-test, *** p<0.001. (G) Quantification of percentage of Sftpc⁺ alveolar organoids, AcTub^+^ airway organoids, double-positive (DP, SftpC^+^/AcTub^+^) and double-negative (DN, SftpC^-^/AcTub^-^) organoids. Data represent mean ± SEM of n = 9 mice. Statistical analysis was performed using two-way ANOVA followed by Sidak’s multiple comparisons test, * indicates differences in SftpC^+^, # indicates differences in DN. ***/### p< 0.001. (H) Representative images of organoid formation after replating, with or without the indicated secondary inflammatory stimuli. Scale bar 500 µm. (I) Quantification of organoid-forming efficiency after replating with or without the second hit stimuli. Data represent mean ± SEM of n = 6-12 mice. Statistical analysis was performed using two-way ANOVA followed by Sidak’s multiple comparisons test, * p<0.05, ** p<0.01, *** p<0.001. (j) Quantification of organoid size after replating under the indicated conditions. Data represent median ± quartiles of n = 6-12 mice. Statistical analysis was performed using Kruskal-Wallis followed by Dunńs multiple comparisons test, * p<0.05, ** p<0.01, *** p<0.001.

As shown in Fig. 4, treatment with 5% CSE significantly reduced organoid formation (Fig. 4B, C) and alveolar differentiation (Fig. 4B, F, G) while increasing organoid size (Fig. 4D), a feature typically associated with impaired alveolar specification, as alveolar organoids are generally smaller. In contrast, airway epithelial differentiation remained unaffected (Fig. 4E, G).

Interestingly, after replating, cells pre-exposed to CSE displayed a markedly reduced capacity to regenerate and form new organoids (Fig. 4H, I). Although the addition of a secondary inflammatory hit increased organoid formation (Fig. 4H, I), consistent with earlier findings. However, the total number of organoids remained significantly lower than in untreated controls (Fig. 4I). Organoids pre-exposed to CSE were less sensitive to IFNγ treatment in terms of organoid numbers (Fig. 3I), mirroring the response seen in organoids derived from *in vivo* CS-exposed mice. However, IFNγ addition produced an opposite effect on organoid size. While it reduced organoid size in organoids derived from *in vivo* CS-exposed mice (Fig. 3B), it increased organoid size in CSE-pre-exposed organoids *in vitro* (Fig. 4J). Staining for SftpC⁺ cell differentiation could not be performed in this second-hit organoid model due to incompatibility with the assay conditions.

Altogether, these results suggest that CSE exposure alters epithelial cell behavior by reducing their proliferative capacity, while simultaneously turning them more resistant to IFNγ-mediated suppression.

### cGAS-STING pathway inhibitors partially reverse cigarette smoke-induced epithelial dysfunction

Given the strong upregulation of interferon-related pathways and cGAS-STING components in our transcriptomic dataset, we investigated whether pharmacological inhibition of this pathway could mitigate the detrimental effects of CS and thereby enhance the regenerative capacity of epithelial progenitors. To this end, we treated lung organoids with a selective cGAS inhibitor: RU.521, in two different contexts: (1) organoids concomitantly exposed to 5% CSE and the drug *in vitro* for 14 days (Fig. 5), and (2) organoids derived from *in vivo* CS-exposed mice (Suppl. Fig. 5).

**Figure 5:**
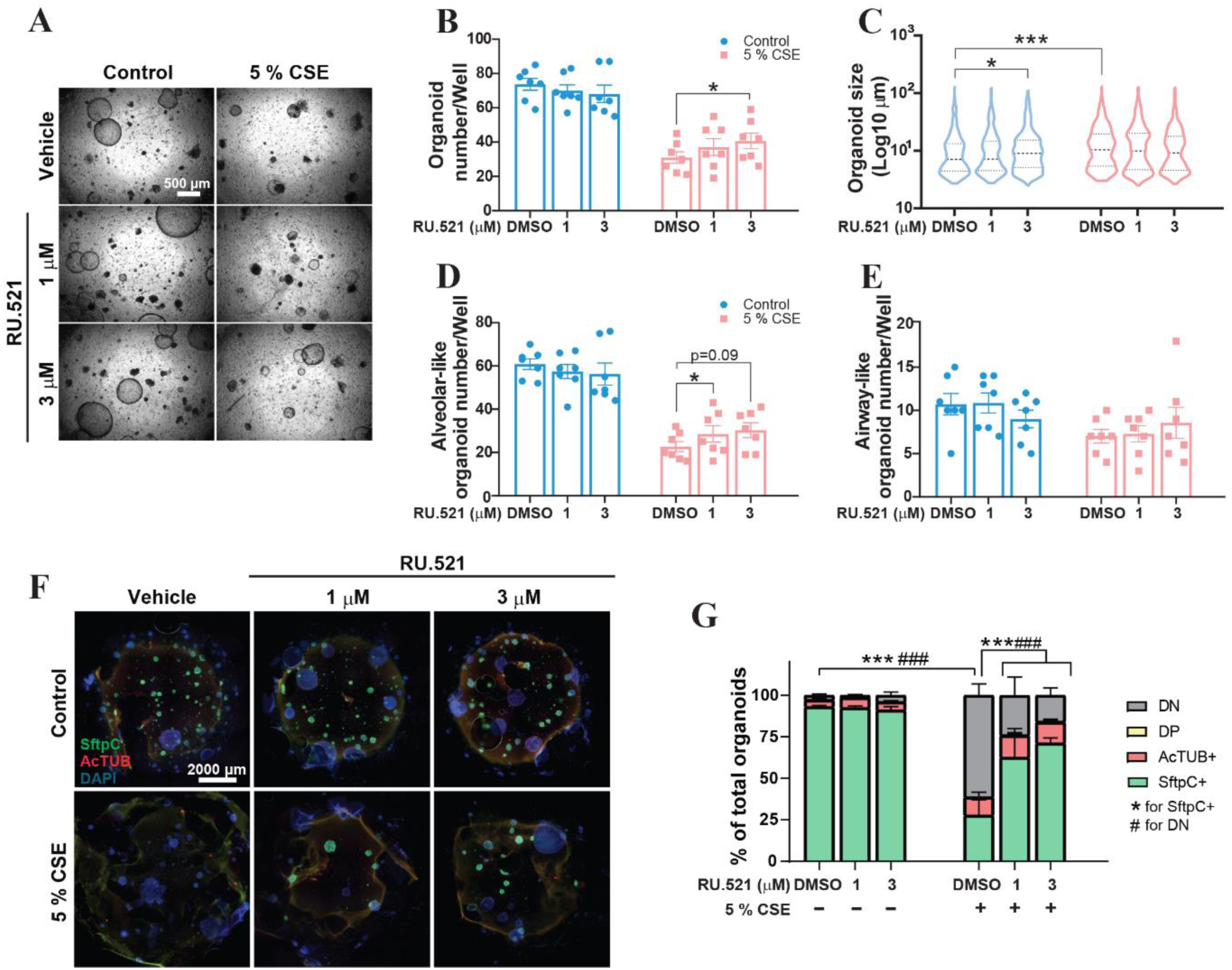
Pharmacological inhibition of cGAS selectively enhances alveolar regeneration in cigarette smoke *in vitro*-exposed organoids. (A) Representative light microscopy images of organoids derived from control and *in vitro* 5 % CSE subjected to increasing concentrations of RU.521 cGAS inhibitor or vehicle (DMSO) (Scale bar 500 µm). (B) Quantification of organoid-forming efficiency. Data represent mean ± SEM of n = 7 mice. Statistical analysis was performed using two-way ANOVA followed by Sidak’s multiple comparisons test, *p<0.05. (C) Quantification of organoid size under the indicated conditions. Data represent median ± quartiles of n = 7 mice. Statistical analysis was performed using Kruskal-Wallis followed by Dunńs multiple comparisons test, *p<0.05, *** p<0.001. Quantification of total airway-like (D) and alveolar-like (E) organoids under the indicated conditions. Data represent mean ± SEM of n = 7 mice. Statistical analysis was performed using two-way ANOVA followed by Sidak’s multiple comparisons test *p<0.05. (F) Representative immunofluorescence images of organoids under the indicated conditions with SftpC^+^ (green) and AcTub+ (red) markers, counterstained with DAPI (blue). (Scale bar 2000 µm). (G) Quantification of percentage of Sftpc⁺ alveolar organoids, AcTub^+^ airway organoids, double-positive (DP, SftpC^+^/AcTub^+^) and double-negative (DN, SftpC^-^/AcTub^-^) organoids. Data represent mean ± SEM of n = 9 mice. Statistical analysis was performed using two-way ANOVA followed by Dunnett’s multiple comparisons test, * indicates differences in SftpC^+^, # indicates differences in DN. ***/### p< 0.001.

Pharmacological inhibition of cGAS using RU.521 significantly increased organoid formation in both the *in vitro* CSE model (Fig. 5A, B) and organoids derived from in vivo CS-exposed mice (Suppl. Fig. 5A, B), whereas no effect was observed in untreated organoids (Fig. 5A, B) or organoids derived from AIR-exposed mice (Suppl. Fig. 5A, B). RU.521 also increased organoid size in untreated control conditions but had no additional effect on size in the 5% CSE-treated groups, where size was already elevated due to CSE exposure (Fig. 5C). A similar trend was observed in organoids derived from *in vivo* CS-exposed mice (Suppl. Fig. 5C), although the RU.521-induced size increase was not statistically significant in the AIR condition.

Notably, RU.521 significantly enhanced the number of alveolar-like organoids (Fig. 5D, & Suppl. Fig. 5D), without altering the number of airway-like organoids (Fig. 5E, & Suppl. Fig. 5E), suggesting a selective improvement in alveolar differentiation. Further immunofluorescence analysis confirmed these findings, showing more SftpC⁺ organoids in both *in vitro* and *in vivo* smoke exposure + RU.521 conditions, whereas no changes were observed in control samples not exposed to CSE (Fig. 5F, G, & Suppl. Fig. 5F, G). This increase was accompanied by a decrease in double-negative (DN) organoids, while the airway (AcTub⁺) organoids remained unchanged across all conditions (Fig. 5F, G, & Suppl. Fig. 5F, G).

Together, these results indicate that pharmacological inhibition of cGAS with RU.521 selectively promotes alveolar regeneration in the context of cigarette smoke exposure, enhancing both organoid formation and differentiation.

### cGAS inhibition with RU.521 enhances epithelial proliferation signatures

To further investigate the *in vitro* effects of RU.521, organoids were cultured in the presence of 5% CSE and RU.521 for 14 days. Organoids were then dissociated and separated into Epcam⁺ epithelial cells and Epcam⁻ cells, the latter enriched in CCL-206 supportive fibroblasts. Normalized counts of epithelial and fibroblast markers further corroborated the enrichment of epithelial markers in the Epcam⁺ fraction (Suppl. Fig. 6A) and fibroblast markers in the CCL-206-enriched Epcam⁻ fraction (Suppl. Fig. 6B), confirming an efficient separation. RNA-seq analysis of these fractions revealed a pronounced transcriptional response to RU.521 in the epithelial compartment, with 1.744 DEGS genes (Fig. 6A). In contrast, a markedly smaller transcriptional response was observed in the fibroblast-enriched fraction, with 364 DEGS (Fig. 6B).

**Figure 6:**
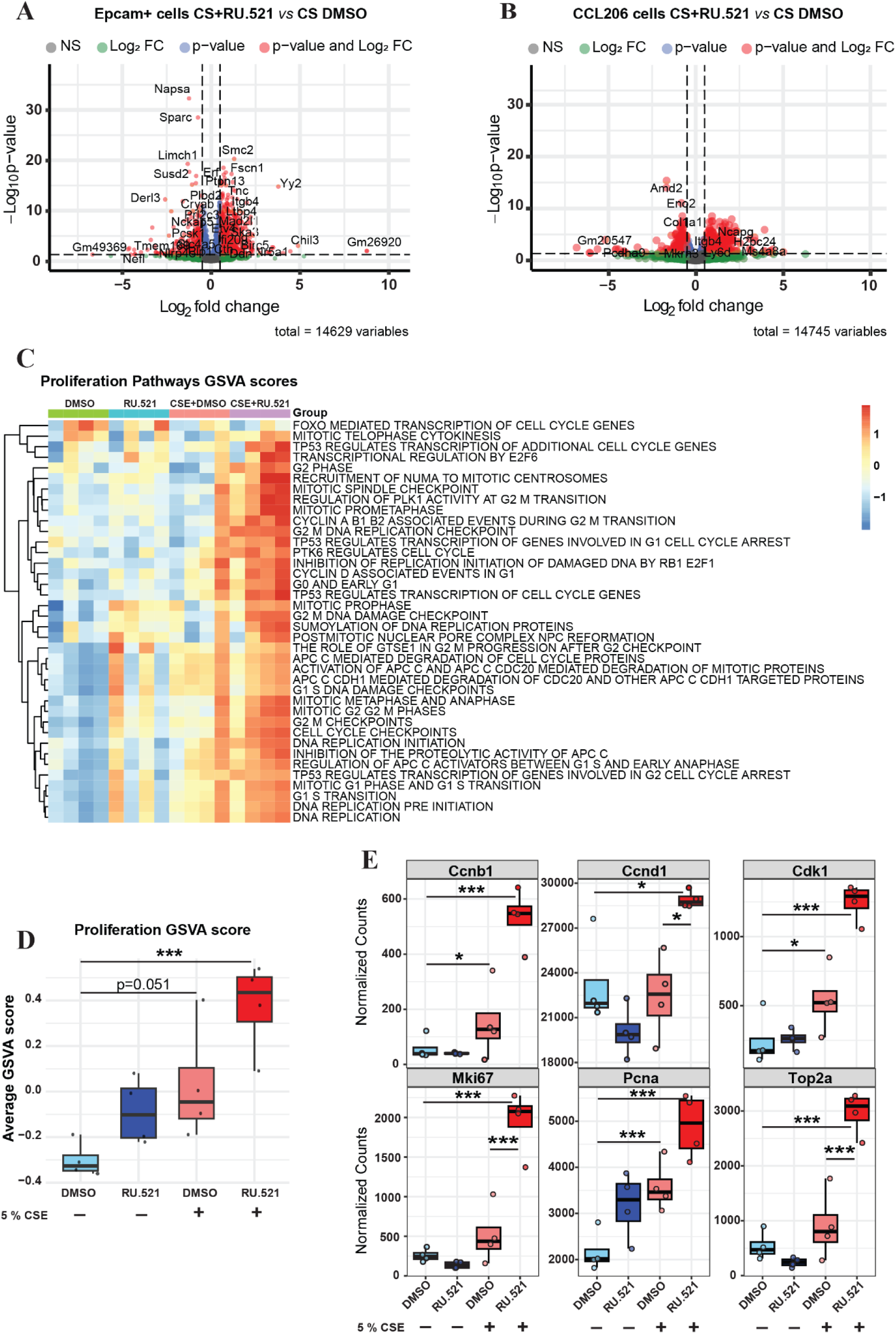
cGAS inhibition with RU.521 induces epithelial proliferation programs in CSE-exposed organoids. (A) Volcano plot of differentially expressed genes (DEGs; padj < 0.05) of a total of 14.629 genes in Epcam⁺ cells. DEGS in CS+RU.521 *vs* CS DMSO samples are highlighted in red. (B) Volcano plot of differentially expressed genes (DEGs; padj < 0.05) of a total of 14.745 genes in CCL-206 fibroblasts. DEGS in CS+RU.521 *vs* CS DMSO samples are highlighted in red. (C) Heatmap showing gene set variation analysis (GSVA) enrichment scores for proliferation-related pathways (Reactome) across Epcam+ samples and conditions. (D) Boxplot summarizing the overall proliferation signature, calculated as the average GSVA enrichment score across all proliferation-related pathways per sample. Data is represented as median with interquartile range and minimum-to-maximum values of n = 4 mice per group. GSVA-derived pathway scores were analyzed using linear mixed-effects models (lme4). Pairwise comparisons were performed using estimated marginal means with Bonferroni correction. Degrees of freedom were estimated using the Satterthwaite approximation (*via* the lmerTest package). *** p<0.001. (E) Normalized expression of selected proliferation-associated genes (*Ccnb1, Ccnd1, Cdk1, Mki67, Pcna,* and *Top2a*) across Epcam+ cells experimental conditions. Values represent normalized counts. Data is represented as median with interquartile range and minimum-to-maximum values of n = 4 mice per group. Statistical significance was determined by differential expression analysis (DESeq2), * p<0.05, *** p<0.001.

In epithelial cells, treatment with the cGAS inhibitor was associated with enrichment of proliferation-related pathways, including cell cycle regulation (*“TP53 regulates transcription of additional cell cycle genes”, “TP53 regulates transcription of cell cycle genes”*), mitotic progression (*”Mitotic prometaphase”, “Mitotic spindle checkpoint”*), and DNA replication (*“DNA replication”*, “*DNA replication preinitiation”*, “*DNA replication initiation*”) (Fig. 6C). This effect was more pronounced in the combined CSE + RU.521 condition (Fig. 6C, D). Consistently, normalized expression of proliferation-associated genes, including *Ccnb1*, *Ccnd1*, *Cdk1*, *Mki67*, *Pcna*, and *Top2a*, was significantly upregulated in the combination treatment. Notably, *Ccnb1*, *Cdk1*, and *Pcna* were also increased in response to CSE alone. These findings are consistent with the increased organoid number and size observed previously (Fig. 5B, C).

To determine whether RU.521 also modulates the supportive function of the mesenchymal compartment, we assessed the expression of key niche factors. Expression levels *of Fgf10, Fgf7, Wnt5a, Wnt2b, Pdgfa,* and *Pdgfb* were not significantly altered upon cGAS inhibition (Suppl. Fig. 7), further suggesting that the primary effects of RU.521 are exerted on the epithelial compartment rather than on mesenchymal support cells.

## Discussion

Chronic lung diseases such as COPD are characterized by a persistent decline in lung function, commonly resulting from chronic exposure to environmental insults like CS or diesel exhaust particles. This decline is further exacerbated by recurrent injury episodes followed by aberrant or incomplete tissue regeneration. Because patients are typically diagnosed only after a measurable drop in lung function, studying the initial events driving defective epithelial repair remains challenging. Therefore, both *in vivo* and *in vitro* models are valuable tools to investigate early pathogenic mechanisms linking chronic inflammation with impaired regeneration of alveolar epithelial progenitors, particularly AT2 cells. In this study, we set out to investigate how chronic CS exposure affects the regenerative capacity of epithelial progenitor cells and their response to secondary inflammatory stimuli. We show that CS induces a sustained interferon-associated transcriptional program in epithelial cells that is linked to impaired alveolar differentiation. Functionally, epithelial progenitors exhibit reduced sensitivity to IFNγ-mediated suppression, indicating a context-dependent reprogramming of their response to inflammatory cues. Furthermore, our data support a role for cGAS-STING signaling in this process, as its pharmacological inhibition partially restores epithelial proliferation and differentiation capacity *in vitro*.

To investigate these processes, we used CS exposure as a model of chronic low-grade inflammation, both *in vivo* and *in vitro*, in combination with a secondary inflammatory stimulus to simulate recurrent injury episodes commonly observed in patients. The 6-week CS exposure protocol led to alterations in lung function consistent with reduced pulmonary elasticity. This aligns with the initial stages of COPD and contrasts with models using prolonged exposure (4–6 months), where emphysematous lesions are already established ^24^. Thus, our model provides a valuable platform to study epithelial changes before irreversible damage occurs and to identify early molecular targets that could promote repair.

With the inclusion of a second hit stimulus in our model, we aimed to mimic the effect of an exacerbation in our *in vitro* organoid system. Unexpectedly, the addition of a secondary inflammatory stimulus on top of an already inflamed environment with cigarette smoke was not detrimental but enhanced epithelial growth, whereas alveolar differentiation was repressed. This effect could be explained by the initial beneficial effect that cytokines have on repair mechanisms as part of the acute inflammation phase, initiating immune responses that clear pathogens and contribute to tissue repair ^25^. On the contrary, treatment with IFNγ as a second hit stimulus was able to significantly impair lung repair in the healthy controls but showed minimal effect on those cells exposed to chronic cigarette smoke. This could be explained by the increased sensitivity of alveolar type II cells that are prone to undergo repair mechanisms ^26,27^.

At the epithelial progenitor level, we observed transcriptomic and functional changes in the Epcam⁺ cell population, including the activation of inflammatory pathways and a reduced capacity to differentiate into Sftpc⁺ AT2-like organoids. Among the inflammatory pathways upregulated, interferon (IFN) signaling emerged as a key driver, likely downstream of cGAS-STING activation. CS exposure is known to trigger the release of nuclear or mitochondrial DNA into the cytosol, activating the cGAS-STING pathway and promoting chronic inflammation and tissue damage ^28,29^. While cGAS-STING plays a protective role during acute damage or infection, its persistent activation may contribute to COPD progression by sustaining inflammation and impairing regenerative responses ^30,31^.

This reprogramming of epithelial cells could explain our findings in the organoid assays. Specifically, Epcam⁺ cells from CS-exposed mice showed a blunted sensitivity to IFNγ in terms of organoid-forming efficiency, compared to control cells, yet their impaired ability to differentiate into alveolar organoids remained unchanged. These results suggest that chronic inflammation either reprograms epithelial progenitors or selects for subpopulations more resistant to IFN-induced suppression. This aligns with recent evidence in human lung tissue, where a subset of IFN-γ–responsive proliferative alveolar epithelial cells was selectively lost in emphysema patients ^16^. Notably, similar effects were observed in both *in vivo* and *in vitro* CS exposure models, supporting the idea that CS itself, independently of immune cues, drives this functional shift. This may reflect a selective enrichment for stress-resistant epithelial subpopulations following chronic inflammatory insult.

The literature on the role of IFNγ in alveolar regeneration remains contradictory. Some studies show that IFNγ disrupts repair by enhancing epithelial cell death ^32^, while others report cell-type-specific effects: negative impacts on AT1 cells but regenerative responses in AT2 cells at low doses ^17^, as well as increased AT2 to AT1 differentiation ^33^. These discrepancies likely reflect the complex interplay between dose, duration, and cellular context. Acute, low-level IFNγ may transiently support repair, while chronic or high-dose exposure impairs it. Interestingly, IFNγ levels in COPD patients remain in the picogram range ^34,35^, and although these levels increase during exacerbations ^35^, they are still lower than those used in our study. Nonetheless, they correlate with impaired lung function and epithelial disruption in patients ^36^, as well as emphysema development in murine models ^37^.

Our results highlight the context-dependent potential of targeting the cGAS-STING pathway to improve epithelial regeneration after cigarette smoke exposure. We demonstrate that pharmacological blockade with the selective cGAS antagonist RU.521 can improve lung repair capacity, increase epithelial proliferation, and promote AT2 cell differentiation *in vitro*, supporting the concept that dampening aberrant immune activation may aid epithelial recovery in COPD. This effect is consistent with growing evidence linking cigarette smoke and particulate matter to double-strand DNA release, thereby further activating the cGAS-STING-IFN axis and perpetuating the chronic inflammation in the lung ^28–30^. Supporting this concept, genetic deletion of *Mb21d1* (cGAS) or *Tmem173* (STING) has been shown to markedly attenuate CS–induced pulmonary inflammation *in vivo*, resulting in reduced neutrophilic infiltration, inflammatory chemokine production, and type I interferon signaling ^29^. In contrast, TLR9 deficiency did not confer similar protection, highlighting the specific contribution of cytosolic DNA sensing through the cGAS-STING pathway in smoke-induced lung injury ^29^.

Importantly, our transcriptomic analysis of RU.521-treated organoids further provides mechanistic insight into these effects, revealing a marked induction of epithelial proliferation programs. The enrichment of cell cycle–related pathways, together with the upregulation of key proliferation markers, suggests that cGAS inhibition primarily promotes expansion of the epithelial progenitor pool. In contrast, mesenchymal niche factor expression remained relatively unchanged, indicating that this effect is predominantly epithelial-intrinsic. These findings provide additional insight into our understanding of how cGAS-STING signaling impacts epithelial repair, suggesting that its inhibition may shift the balance of progenitor cell states from a dysfunctional or stress-associated program toward a more proliferative and regenerative phenotype.

Inhibition of this pathway has, therefore, attracted considerable interest across disease models. H-151, a small molecule inhibiting STING, has proven to reduce lung inflammation and fibrosis in models of idiopathic pulmonary fibrosis (IPF) ^38^ and acute lung injury ^39^. Similarly, RU.521 has shown efficacy in ameliorating inflammatory cytokine release other inflammatory disease models, such as imiquimod psoriasis-like dermatitis ^40^ and ISD017 in lupus erythematosus ^41,42^. Several orally available cGAS inhibitors have recently entered clinical development, including VENT-03 (Ventus Therapeutics) and IMSB301 (ImmuneSensor Therapeutics), both of which have reported favorable phase 1 safety and pharmacodynamic data in company communications, although peer-reviewed clinical datasets are not yet available.

Although direct evidence in COPD remains limited, our data suggest that excessive cGAS-STING signaling may represent a driver in impaired epithelial repair. Therapeutic targeting of this pathway could therefore not only reduce chronic interferon type I signaling but also restore the regenerative potential of AT2 progenitors. In this regard, future studies are warranted to explore the translational potential of cGAS-STING inhibition not only in IPF or autoimmune diseases, but also in COPD.

## Author contributions

R.F.M., B.N.M., and R.G. designed the study. R.F.M., P.A.S., V.V., I.C.G., and J.C.W., performed experiments. R.F.M., P.A.S., I.S.T.B. and J.C.W. analyzed data. The manuscript was drafted by R.F.M. and revised by R.G. and B.N.M. All authors participated in manuscript preparation and provided final approval of the submitted work.

## Acknowledgements

We thank D.F.N. for assistance with the deconvolution analysis presented in Supplementary Figure 1A.

## Data Availability

Raw RNA-seq FASTQ files together with processed count matrices have been deposited in the Gene Expression Omnibus (GEO) database (https://www.ncbi.nlm.nih.gov/geo) under accession number GSE334950.

## Clinical relevance

Impaired epithelial repair is a key feature of COPD and is closely linked to chronic inflammation. Here, we show that cigarette smoke exposure alters interferon signaling and the regenerative capacity of alveolar epithelial cells. Our findings suggest that modulation of the cGAS-STING pathway may help to restore epithelial repair responses under chronic inflammatory conditions.

## References

1. Hogg JC, Senior RM. Chronic obstructive pulmonary disease • 2: Pathology and biochemistry of emphysema. Thorax. Published online 2002. doi:10.1136/thorax.57.9.830

2. Hogg JC, Timens W. The pathology of chronic obstructive pulmonary disease. Annu Rev Pathol Mech Dis. Published online 2009. doi:10.1146/annurev.pathol.4.110807.092145

3. Fischer BM, Voynow JA, Ghio AJ. COPD: Balancing oxidants and antioxidants. Int J COPD. Published online 2015. doi:10.2147/COPD.S42414

4. Wang Y, Xu J, Meng Y, Adcock IM, Yao X. Role of inflammatory cells in airway remodeling in COPD. Int J COPD. Published online 2018. doi:10.2147/COPD.S176122

5. Barnes PJ. Inflammatory mechanisms in patients with chronic obstructive pulmonary disease. J Allergy Clin Immunol. Published online 2016. doi:10.1016/j.jaci.2016.05.011

6. Boukhenouna S, Wilson MA, Bahmed K, Kosmider B. Reactive oxygen species in chronic obstructive pulmonary disease. Oxid Med Cell Longev. Published online 2018. doi:10.1155/2018/5730395

7. Brightling C, Greening N. Airway inflammation in COPD: Progress to precision medicine. Eur Respir J. Published online 2019. doi:10.1183/13993003.00651-2019

8. Song L, Li K, Chen H, Xie L. Cell Cross-Talk in Alveolar Microenvironment: From Lung Injury to Fibrosis. Am J Respir Cell Mol Biol. 2024;71(1):30–42. doi:10.1165/rcmb.2023-0426TR

9. Devi K, Parekh KR. Mechanotransduction: A Master Regulator of Alveolar Cell Fate Determination. Bioengineering. 2025;12(7):760. doi:10.3390/bioengineering12070760

10. Zhang J, Liu Y. Epithelial stem cells and niches in lung alveolar regeneration and diseases. Chinese Med J Pulm Crit Care Med. 2024;2(1):17–26. doi:10.1016/j.pccm.2023.10.007

11. Bender EC, Tareq HS, Suggs LJ. Inflammation: a matter of immune cell life and death. npj Biomed Innov. 2025;2(1):7-. doi:10.1038/s44385-025-00010-4

12. Chavda VP, Feehan J, Apostolopoulos V. Inflammation: The Cause of All Diseases. Cells. 2024;13(22):1906. doi:10.3390/cells13221906

13. Mulvanny A, Pattwell C, Beech A, Southworth T, Singh D. Validation of Sputum Biomarker Immunoassays and Cytokine Expression Profiles in COPD. Biomedicines. 2022;10(8). doi:10.3390/biomedicines10081949

14. Kortekaas RK, Geillinger-Kästle KE, Fuentes-Mateos R, et al. The disruptive effects of COPD exacerbation-associated factors on epithelial repair responses. Front Immunol. 2024;15:1346491. doi:10.3389/FIMMU.2024.1346491/BIBTEX

15. Ciminieri C, Woest ME, Reynaert NL, et al. IL-1b Induces a Proinflammatory Fibroblast Microenvironment that Impairs Lung Progenitors’ Function. Am J Respir Cell Mol Biol. 2023;68(4):444–455. doi:10.1165/rcmb.2022-0209OC

16. Wang C, Hyams B, Allen NC, et al. Dysregulated lung stroma drives emphysema exacerbation by potentiating resident lymphocytes to suppress an epithelial stem cell reservoir. Immunity. 2023;56(3):576–591.e10. doi:10.1016/J.IMMUNI.2023.01.032

17. Dost AFM, Balážová K, Pou Casellas C, et al. Interferon-γ selectively promotes survival of alveolar progenitor cells in a human lung organoid model. EMBO J. 2026;45(10):3364–3395. doi:10.1038/s44318-026-00774-4

18. Wu X, Sophie I, Conlon TM, et al. A transcriptomics-guided drug target discovery strategy identifies receptor ligands for lung regeneration. Sci Adv. 2022;8(12):9949. doi:10.1126/sciadv.abj9949

19. van der Koog L, Woest ME, Gorter IC, et al. Fibroblast-derived osteoglycin promotes epithelial cell repair. npj Regen Med. 2025;10(1):16. doi:10.1038/s41536-025-00404-3

20. Kortekaas RK, Geillinger-Kästle KE, Fuentes-Mateos R, et al. The soluble factor milieu in idiopathic pulmonary fibrosis dysregulates epithelial differentiation. FASEB J. 2024;38(19):e70077. doi:10.1096/fj.202302405RR

21. Willems SH, Qian S, Lång P, et al. TRAPping the effects of tobacco smoking: the regulation and function of Acp5 expression in lung macrophages. 101152/ajplung001572024. 2025;328(4):L497–L511. doi:10.1152/AJPLUNG.00157.2024

22. Sabogal-Guáqueta AM, Marmolejo-Garza A, Trombetta-Lima M, et al. Species-specific metabolic reprogramming in human and mouse microglia during inflammatory pathway induction. Nat Commun. 2023;14(1). doi:10.1038/s41467-023-42096-7

23. Angelidis I, Simon LM, Fernandez IE, et al. An atlas of the aging lung mapped by single cell transcriptomics and deep tissue proteomics. Nat Commun 2019 101. 2019;10(1):1-17. doi:10.1038/s41467-019-08831-9

24. John-Schuster G, Hager K, Conlon TM, et al. Cigarette smoke-induced iBALT mediates macrophage activation in a B cell-dependent manner in COPD. Am J Physiol - Lung Cell Mol Physiol. 2014;307(9):L692–L706. doi:10.1152/AJPLUNG.00092.2014/ASSET/IMAGES/LARGE/ZH50201466050008.JPEG

25. Yamada M, Fujino N, Ichinose M. Inflammatory responses in the initiation of lung repair and regeneration: Their role in stimulating lung resident stem cells. Inflamm Regen. 2016;36(1):15-. doi:10.1186/s41232-016-0020-7

26. Okutomo K, Fujino N, Yamada M, et al. Increased LHX9 expression in alveolar epithelial type 2 cells of patients with chronic obstructive pulmonary disease. Respir Investig. 2022;60(1):119–128. doi:10.1016/j.resinv.2021.08.007

27. Theron M, Huang KJ, Chen YW, Liu CC, Lei HY. A probable role for IFN-γ in the development of a lung immunopathology in SARS. Cytokine. 2005;32(1):30–38. doi:10.1016/j.cyto.2005.07.007

28. Tian M, Li F, Pei H, Liu X, Nie H. The role of the cGAS-STING pathway in chronic pulmonary inflammatory diseases. Front Med. 2024;11. doi:10.3389/FMED.2024.1436091,

29. Nascimento M, Gombault A, Lacerda-Queiroz N, et al. Self-DNA release and STING-dependent sensing drives inflammation to cigarette smoke in mice. Sci Reports 2019 91. 2019;9(1):1-10. doi:10.1038/s41598-019-51427-y

30. Liao K, Wang F, Xia C, et al. The cGAS-STING pathway in COPD: targeting its role and therapeutic potential. Respir Res. 2024;25(1):302. doi:10.1186/s12931-024-02915-x

31. Zhang Z, Jiang J, Wu G, Wei X, Weng Y, Huang LS. The cGAS-STING Pathway in Pulmonary Diseases: Mechanisms and Therapeutic Potential. Int J Mol Sci. 2025;26(21):10423. doi:10.3390/ijms262110423

32. Katsura H, Sontake V, Tata A, et al. Human Lung Stem Cell-Based Alveolospheres Provide Insights into SARS-CoV-2-Mediated Interferon Responses and Pneumocyte Dysfunction. Cell Stem Cell. 2020;27(6):890–904.e8. doi:10.1016/j.stem.2020.10.005

33. Zhang X, Ali M, Pantuck MA, et al. CD8 T cell response and its released cytokine IFN-γ are necessary for lung alveolar epithelial repair during bacterial pneumonia. Front Immunol. 2023;14:1268078. doi:10.3389/fimmu.2023.1268078

34. Moermans C, Heinen V, Nguyen M, et al. Local and systemic cellular inflammation and cytokine release in chronic obstructive pulmonary disease. Cytokine. 2011;56(2):298–304. doi:10.1016/j.cyto.2011.07.010

35. Budroni S, Taccone M, Stella M, et al. Cytokine Biomarkers of Exacerbations in Sputum From Patients With Chronic Obstructive Pulmonary Disease: A Prospective Cohort Study. J Infect Dis. 2024;230(5):e1112–e1120. doi:10.1093/INFDIS/JIAE232,

36. Kapellos TS, Conlon TM, Yildirim AÖ, Lehmann M. The impact of the immune system on lung injury and regeneration in COPD. Eur Respir J. 2023;62(4). doi:10.1183/13993003.00589-2023

37. Wang Z, Zheng T, Zhu Z, et al. Interferon Induction of Pulmonary Emphysema in the Adult Murine Lung. J Exp Med. 2000;192(11):1587–1599. Accessed February 4, 2025. http://www.jem.org/cgi/content/full/192/11/1587

38. Zhang J, Du J, Liu D, et al. Polystyrene microplastics induce pulmonary fibrosis by promoting alveolar epithelial cell ferroptosis through cGas/STING signaling. Ecotoxicol Environ Saf. 2024;277:116357. doi:10.1016/j.ecoenv.2024.116357

39. Ge X, Liu Q, Fan H, et al. STING facilitates the development of radiation-induced lung injury via regulating the PERK/eIF2α pathway. Transl Lung Cancer Res. 2024;13(11):3010–3025. doi:10.21037/tlcr-24-649

40. Zhang Z, Wei X, Huang Q, et al. Discovery of STING antagonists targeting cGAS-STING pathway to alleviate IMQ-induced psoriasis-like dermatitis. Eur J Pharm Sci. 2025;210. doi:10.1016/j.ejps.2025.107091

41. Alee I, Chantawichitwong P, Leelahavanichkul A, Paludan SR, Pisitkun T, Pisitkun P. The STING inhibitor (ISD-017) reduces glomerulonephritis in 129.B6.Fcgr2b-deficient mice. Sci Rep. 2024;14(1). doi:10.1038/s41598-024-61597-z

42. Prabakaran T, Troldborg A, Kumpunya S, et al. A STING antagonist modulating the interaction with STIM1 blocks ER-to-Golgi trafficking and inhibits lupus pathology. EBioMedicine. 2021;66:103314. doi:10.1016/j.ebiom.2021.103314

