## Supplementary material for "cGAS-STING drives alveolar epithelial cell dysfunction in cigarette smoke-induced lung injury"

**Supplementary Materials and Methods**

*Animal handling*

Mouse experiments were performed at the central animal facility (CDP) of the University Medical Center of Groningen (UMCG) following the national guidelines and upon approval of the experimental procedures by the CDP and the institutional animal care and use committee of the University of Groningen (license AVD10500209120). C57Bl6 male mice aged 8-12 weeks were housed under a 12-hour light/dark cycle and allowed food and water *ad libitum*.

*Chronic in vivo cigarette smoke exposure*

Mice were randomly divided into two groups (n= 8 per group): fresh air (AIR) and cigarette smoke exposure (CS). The number of animals per group was determined based on a power analysis using data from an acute cigarette smoke exposure previously performed in our lab[1]. In that study, mice exposed to air generated an average of 303 ± 55 alveolar organoids per animal, while those exposed to cigarette smoke for one week generated approximately 230 ± 46, Based on a two-tailed t-test with $\alpha = 0.05$ and a power ($1-\beta$) of 0.8, this indicated that 8 animals per group would be sufficient to detect this difference.

Mice were whole-body exposed to 3R4F research cigarettes (Tobacco Research Institute, University of Kentucky, Lexington, KY), as previously described in [1], twice a day for 6 weeks to establish chronic smoke-induced inflammation. The animals were initially exposed to 1 cigarette in the morning and 3 in the afternoon on day 1, and from day 2 to 42, mice were exposed to 5 cigarettes in each session. All cigarettes were smoked without a filter for 5 minutes at a rate of 5L/h in a ratio of 60L/h air, using a peristaltic pump (46 rpm, Watson Marlow 323 E/D, Rotterdam, NL). In the control group, mice were exposed to fresh air using similar chambers to those of the CS group. 16 hours after the last smoking session, mice were subjected to lung function experiments and subsequently terminated for lung tissue extraction.

*Lung function measurements*

Lung function measurements were performed as previously described in [2]. Briefly, mice were anesthetized with a combination of 0.5 mg/kg Dexdormitor® and 40 mg/kg Ketamine. After anesthesia was achieved, rocuronium bromide (Fresenius Kabi, 10 mg/mL) was administered intraperitoneally (i.p.) to relax the muscles. Mice were then subjected to a tracheotomy to insert a cannula in the trachea, and they were subsequently connected to the FlexiVent system module 2 (Scireq). The animals were allowed to adjust to the ventilator for 2 minutes of mechanical ventilation with a tidal volume of 10 mL/kg at a frequency of 150 breaths/min, and lung function parameters were assessed using a pre-installed protocol for SnapShot, Primewave perturbation, and forced expired volume maneuver using the Flexiware V8.3.0 software. Per mouse, 3 individual measurements were acquired.

*Tissue processing*

Lung lobes were fixed overnight at constant pressure by intratracheal instillation of 10% (v/v) neutral-buffered formalin (J.T. Baker, 3933.9020). Tissues were subsequently embedded in paraffin (Epredia, 6774060) and sectioned at 5 µm thickness. For histological analysis, sections were stained with hematoxylin and eosin (H&E). Whole-slide images were acquired using a Hamamatsu NanoZoomer 2.0HT digital slide scanner at 40× magnification. Airspace enlargement was quantified by measuring the mean linear intercept (MLI) using a semi-automated, unbiased morphometric method, analyzing 10 fields of view per lung [3].

*Murine primary lung epithelial and macrophages isolation*

Primary epithelial progenitor cells (CD45-/CD31-/Epcam+ cells) were isolated as described previously in [2]. Briefly, lung lobes were digested in dispase (#354235, Corning) at room temperature (RT) for 45 min, followed by mechanical disruption of the tissue. The resulting suspension was passed through a 100 µm cell strainer, incubated with CD45 (Miltenyi Biotec, Teterow, Germany #130–052–301) and CD31 (Miltenyi, #130–097–418) microbeads for 20 min at 4 °C. After depletion of CD45+/CD31+ fraction, the CD45-/CD31- cell fraction was subsequently enriched for epithelial cells by positive selection using the Epcam (CD326) microbeads (Miltenyi #130–105–958). Epcam+ cells were resuspended in DMEM/F-12 with 10 % FBS. Prior findings based on a combination of RNAseq, flow cytometry data, and immunofluorescence stainings indicate this cell population consists of ~80% AT2 cells [4,5]. Macrophages were isolated from the CD31+/CD45+ fraction as previously described in [6]. Briefly, CD31+/CD45+ cell fraction was incubated in 6-well plates for 30 min at 37 °C in RPMI-1640 medium (Gibco, Bleiswijk, The Netherlands) supplemented with 10 % FBS and 10 µg/mL gentamycin. Nonadherent cells were then washed away, and macrophage cells were subsequently stored at −80 °C.

*CCL-206 murine lung fibroblast cell culture*

Mouse fibroblasts, CCL-206 [Mlg (CCL-206); American Type Culture Collection (ATCC), Wesel, Germany] were cultured in a combination (1:1) of Dulbecco's modified Eagle's medium (DMEM)/F12 medium supplemented with 10% (v/v) fetal bovine serum (FBS), penicillin/streptomycin (100 U/ml), 2 mM L-glutamine, and 1% amphotericin B, in a humidified atmosphere under 5% CO2/95% air at 37 °C. For organoid experiments, fibroblasts were proliferation-inactivated by incubation in mitomycin C (10 µg/ml; M4287, Sigma-Aldrich) for

2 hours, followed by three washes with phosphate-buffered saline (PBS), after which the cells were trypsinized for the organoid co-cultures.

*Organoid culture*

The organoid culture system was based on previously published protocols [2,4,5]. Briefly, $10^4$ freshly isolated epithelial progenitor cells (CD45-/CD31-/Epcam+) were co-cultured with $10^4$ proliferation-inactivated CCL-206 in a mixture consisting of 60% non-growth factor reduced Matrigel® (#11543550, Fisher Scientific, Landsmeer, NL) and 40 % lung fibroblast medium. This mixture (total of 100 µl) was added to a 24-well culture insert (Corning, USA) in a 24-well plate containing 420 µl of organoid media [DMEM/F-12 supplemented with 5% FBS, 100 U/ml penicillin/streptomycin, 2 mM L-glutamine, 1 % amphotericin B, 0.025 µg/ml murine epidermal growth factor (EGF), 1% insulin-transferrin-selenium, and 30 µg/ml bovine pituitary extract]. Y-27632 Rock inhibitor (10 µM, #1254, Tocris Bioscience, Oxford, UK) was added for the first 48 h of culture. Organoids were treated with the compounds listed in **Table 1** during the 14 days of culture, and the medium was replaced every 2-3 days. After 14 days of culture, organoid size (diameter) and number were analyzed through brightfield images and Fiji ImageJ software (NIH).

*Cigarette smoke extract*

Cigarette smoke extract (CSE) was prepared as described in [1,2]. Briefly, the smoke of two 3R4F research cigarettes was pumped into 25 mL of warm 1:1 DMEM/F12 supplemented with 10% FBS and PS to produce 100% CSE. All cigarettes were smoked without a filter, and smoke passed through the medium using a peristaltic pump (45 rpm, Watson Marlow 323 E/D, Rotterdam, The Netherlands). CSE was freshly prepared before each media change and further diluted to a 5% working concentration for the organoid assays.

*Two-hit organoid culture*

The organoid culture was modified to allow a two-hit exposure *in vitro* organoid model. The first hit consisted of treating the organoids for 14 days with 5% cigarette smoke extract (CSE). The second hit involved exposure to the compounds stated in **Table 1**. Murine Epcam+ cells were isolated from wild-type mice as stated in the previous section, and co-cultured with CCL-206 in Matrigel® for 14 days in the presence or absence of 5% CSE. After 14 days, Matrigel® was dissolved by adding 100 µl of dispase (#354235, Corning) for 45 min at 37 ºC. The reaction was stopped by adding 200 µl of 5% BSA MACs buffer, organoids were collected into 1.5 mL Eppendorf cups and centrifuged for 5 min at 300 g. Organoid pellets were resuspended in 500 µl TrypLE™ (#12604013, Gibco) and incubated for 15 min at 37 ºC to dissociate them into a cell suspension. The reaction was stopped by adding 800 µl of 10% FBS DMEM/F-12 media, after which cells were centrifuged for 5 min at 300 g and counted. $10^4$ of these cells were co-cultured with $10^4$ proliferation-inactivated CCL-206 fibroblasts in Matrigel® as described before, and organoids were cultured for a further 14 days in the presence of a second hit stimulus (**Table 1**). After 14 days of culture, organoid size (diameter), and number were analyzed through brightfield images and ImageJ software (NIH).

*Organoid dissociation and separation of epithelial and mesenchymal fractions*

For organoid re-sorting experiments, a mixture of $3 \times 10^5$ $Epcam^+$ cells and $3 \times 10^5$ proliferation-inactivated CCL-206 fibroblasts was embedded in 1 mL of Matrigel diluted 1:1.5 (v/v) with DMEM/F12 supplemented with 10% FBS and plated in one well of a 6-well plate as previously described in [2]. After Matrigel polymerization (1 h at 37°C), 2 mL of organoid culture medium was added on top, containing either vehicle or 5% CSE in the presence of DMSO or 3 µM RU.521 (InvivoGen, inh-ru521-2).

After 14 days of culture, Matrigel was dissociated by incubation with dispase (Corning, 354235) for 30 min at 37°C. The reaction was stopped by adding MACS buffer (MACS rinsing solution (Miltenyi Biotec, 130-091-222) supplemented with BSA (Miltenyi Biotec, 130-091-376)). Organoids were collected and centrifuged at 300 × g for 5 min.

Pellets were resuspended in 5 mL diluted trypsin (1:5 in PBS, v/v; T7409, Sigma-Aldrich) and incubated for 5 min at 37°C to obtain a single-cell suspension. Trypsin activity was neutralized by adding 9 mL DMEM/F-12 supplemented with 10% FBS, followed by centrifugation at 300 × g for 5 min.

Cells were incubated with Epcam/CD326 microbeads for 20 min, resuspended in MACS buffer, and separated using a QuadroMACS™ Separator system. This allowed the isolation of $CD326^+$ ($Epcam^+$) epithelial cells and $CD326^-$ fibroblast-enriched fractions derived from organoids, which were subsequently used for RNA extraction and RNA-seq analysis.

*RNA extraction and sequencing*

RNA was isolated from freshly isolated Epcam+ cells and macrophages from mice exposed to fresh air or CS for 6 weeks, or from re-sorted organoid experiments, obtaining $Epcam^+$ and $Epcam^-$ (CCL-206) fractions. Cells were washed 3 times with cold PBS and spun down at 300 g for 5 min. RNA was isolated using Maxwell® 16 LEV Simply RNA Cells Kit (AS1270, Promega Corporation, USA) according to the manufacturer's instructions. RNA sequencing was performed by Biomarker Technologies (BMK, Münster, Germany, www.bmkgene.com/de) using an Illumina NovaSeq 6000 sequencer. The procedure included data quality control, adapter trimming, alignment of short reads, and feature counting. Library preparation was validated by calculating ribosomal (and goblin) content. Checks for possible sample and barcode contaminations were performed, and a set of standard quality metrics for the raw data set was determined using quality control tools (FstQC and FastQA). Reads were trimmed for adapter sequences before alignment using Trimmomatic, and sample reads were aligned to the ensemble mouse reference GRCm39.

*RNA sequencing analysis*

Differential expression analysis was performed and visualized using R version 4.2.0 [7]. Differential expression (DE) was obtained by using DESeq2 version 1.42 [8]. Gene Set Enrichment Analysis (GSEA) was performed using the fgsea package version 1.28 [9] with the following parameters: minimum gene set size of 5 and maximum of 500. Pretty Heatmap (pheatmap) version 1.0.12 [10] was used for heatmap visualization with Manhattan clustering. Ggplot2 version 3.5.0 [11] was used for the volcano, correlation, and bar plots visualization.

Correlation analyses were performed in R using the base package stats, with Pearson's correlation coefficients and corresponding p-values obtained via the functions cor() and cor.test(). Gene set variation analysis (GSVA) was conducted using the GSVA package version 1.50 [12], applying Gaussian kernel estimation and restricting pathway sizes to between 5 and 500 genes.

*Discovery-based proteomics analyses*

Proteome analyses were performed on freshly isolated Epcam+ cells. Briefly, $1x10^6$ cells were washed 3 times with cold PBS, spun down at 300 g for 5 min, and resuspended in 25 µL NP40 lysis solution (0.1% IGEPAL, 0.4 M NaCl, 10 mM Tris-HCl (pH 8.0), 1 mM EDTA). The lysates were incubated for 35 minutes on ice, with a vortexing step halfway through the incubation after which the lysates were spun down for 10 minutes at 12000 g and 4 °C. The soluble fraction was pipetted off and mixed with 9 µL 4x LDS loading buffer (NuPAGE), incubated for 10 minutes at 70 ˚C and 25 µL from this was loaded on a precast 4-12% Bis-Tris gels (Thermo

Scientific). In-gel digestion of the lysates and discovery-based mass spectrometric analyses applying label-free quantification (LFQ) were performed as described previously [13], injecting 5% of the sample for the LC-MS measurements. LC-MS raw data were processed with Spectronaut (version 18.4.231011) (Biognosys) using the standard settings of the directDIA workflow except that quantification was performed on MS1, with a reviewed mouse SwissProt database (www.uniprot.org, 17141 entries). For the quantification, local normalization was applied and the Q-value filtering was set to the classic setting without imputing.

The normalized counts were analyzed for differentially regulated proteins with a significance of padj<0.05 and a log2foldchange threshold of 0.5. Data was further visualized with RStudio. Ggplot2 version 3.5.0 was used for volcano plot, correlation plot, and bar plot visualization.

*Immunofluorescence staining*

Immunofluorescence stainings were performed in the whole insert as previously described in [2]. Briefly, after 14 days of culture, organoids were fixed with ice-cold Acetone-Methanol (1:1, v:v) solution for 15 min at -20 ºC. Then, organoids were blocked in 5% BSA, 2% donkey serum, and 0.1% Triton in PBS overnight (o/n) at 4 ºC. Primary antibodies were diluted 1:200 (pro-SftpC, #AB3786, Millipore; Acetylated α-Tubulin (Ac-Tub), #sc-23950, Santa Cruz Biotechnology) in the antibody solution consisting of 2% BSA, 2% donkey serum, and 0.1% Triton in PBS, and incubated 48 h at 4 ºC. Samples were washed 3 times with PBS and incubated with the secondary antibodies (AlexaFluor488 donkey anti-rabbit, #A21206, Invitrogen; AlexaFluor568 donkey anti-mouse, #A10037, Invitrogen ) for 2 h at RT in the antibody solution. After 3 washes with PBS, membranes were detached from the insert and mounted with Fluoroshield mounting media with DAPI (#AB104139, abcam). Whole-membrane images were taken by using the Nikon Tti microscope (check), and organoids were classified as alveolar ($SftpC^+$), airway (Ac-Tub+), double-positive ($SftpC^+$/$Ac\text{-}Tub^+$), or double-negative (SftpC-/Ac-Tub-) using the cell counter plug-in from ImageJ.

*Statistical analysis*

Data are presented as mean ± SEM for normally distributed data or median with interquartile range for non-normally distributed data. Normality was assessed using four tests (D'Agostino–Pearson, Anderson–Darling, Shapiro–Wilk, and Kolmogorov–Smirnov); data were considered normally distributed when at least three tests indicated normality. For comparisons between two groups, unpaired or paired two-tailed Student's t-tests were applied for parametric data, whereas Mann–Whitney or Wilcoxon tests were used for non-parametric data, as appropriate. For multiple group comparisons, one-way ANOVA followed by Dunnett's post hoc or two-way ANOVA followed by Sidak´s post hoc tests were used for parametric data. When required, appropriate non-parametric alternatives were applied. For GSVA analyses, linear mixed-effects models were used to account for repeated measurements, with Group included as a fixed effect and Mouse as a random effect. Post hoc comparisons were performed with multiple testing correction, as specified in the figure legends.

Gene-level comparisons of normalized RNA-seq counts between $Epcam^+$ and $Epcam^-$ fractions were performed using Wilcoxon rank-sum tests, followed by Benjamini–Hochberg correction for multiple testing.

Sample sizes, number of biological replicates, and statistical tests used for each experiment are indicated in the corresponding figure legends. Statistical significance was defined as $p < 0.05$. Analyses were performed using GraphPad Prism (version 10) and R (version 4.2.0), with RNA-seq and proteomics analyses conducted in R.

**References**


1. Wu X, Sophie I, Conlon TM, et al. A transcriptomics-guided drug target discovery strategy identifies receptor ligands for lung regeneration. *Sci Adv*. 2022;8(12):9949. doi:10.1126/sciadv.abj9949

2. van der Koog L, Woest ME, Gorter IC, et al. Fibroblast-derived osteoglycin promotes epithelial cell repair. *npj Regen Med*. 2025;10(1):16. doi:10.1038/s41536-025-00404-3

3. Salaets T, Tack B, Gie A, et al. A semi-automated method for unbiased alveolar morphometry: Validation in a bronchopulmonary dysplasia model. *PLoS One*. 2020;15(9 September 2020):e0239562. doi:10.1371/journal.pone.0239562

4. Kortekaas RK, Geillinger-Kästle KE, Fuentes-Mateos R, et al. The soluble factor milieu in idiopathic pulmonary fibrosis dysregulates epithelial differentiation. *FASEB J*. 2024;38(19):e70077. doi:10.1096/fj.202302405RR

5. Kortekaas RK, Geillinger-Kästle KE, Fuentes-Mateos R, et al. The disruptive effects of COPD exacerbation-associated factors on epithelial repair responses. *Front Immunol*. 2024;15:1346491. doi:10.3389/FIMMU.2024.1346491/BIBTEX

6. Willems SH, Qian S, Lång P, et al. TRAPping the effects of tobacco smoking: the regulation and function of Acp5 expression in lung macrophages. *https://doi.org/101152/ajplung001572024*. 2025;328(4):L497-L511. doi:10.1152/AJPLUNG.00157.2024

7. R Core Team. R: A Language and Environment for Statistical Computing. Published online 2022. https://www.r-project.org/

8. Love MI, Huber W, Anders S. Moderated estimation of fold change and dispersion for

RNA-seq data with DESeq2. *Genome Biol*. 2014;15(12):550-. doi:10.1186/s13059-014-0550-8

9. Korotkevich G, Sukhov V, Sergushichev A. fgsea: Fast Gene Set Enrichment Analysis. *bioRxiv*. Published online February 1, 2023:1-29. doi:10.1101/060012

10. Kolde R. Pretty Heatmaps [R package pheatmap version 1.0.12]. *CRAN Contrib Packag*. Published online June 5, 2019. doi:10.32614/CRAN.PACKAGE.PHEATMAP

11. Wickham H. Data Analysis. Published online 2016:189-201. doi:10.1007/978-3-319-24277-4_9

12. Hänzelmann S, Castelo R, Guinney J. GSVA: Gene set variation analysis for microarray and RNA-Seq data. *BMC Bioinformatics*. 2013;14. doi:10.1186/1471-2105-14-7

13. Sabogal-Guáqueta AM, Marmolejo-Garza A, Trombetta-Lima M, et al. Species-specific metabolic reprogramming in human and mouse microglia during inflammatory pathway induction. *Nat Commun*. 2023;14(1). doi:10.1038/s41467-023-42096-7

**Supplementary figures**

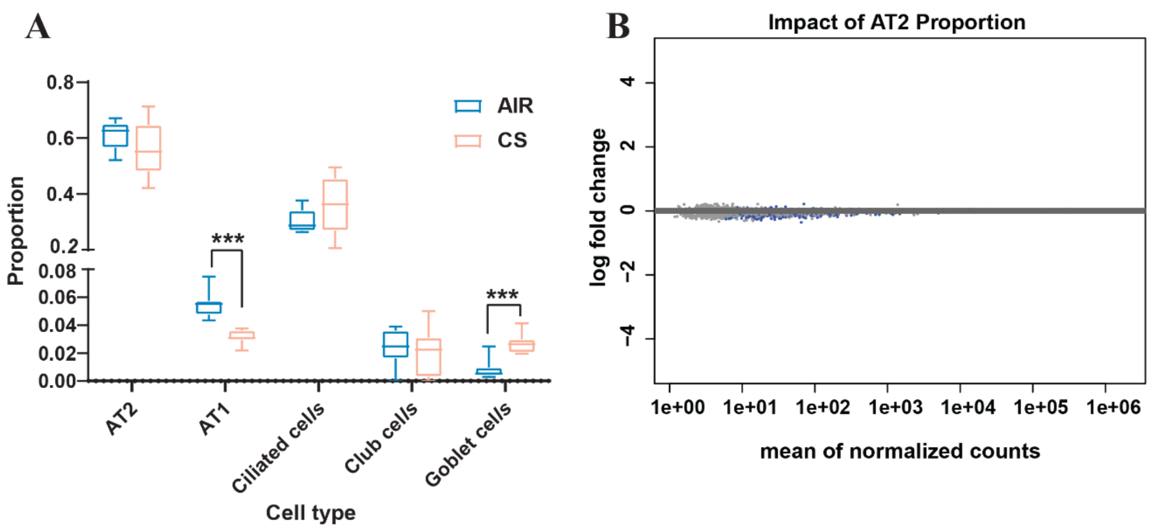


**Supplementary Figure 1: Cell-type deconvolution analysis of isolated Epcam⁺ cells and impact of AT2 proportion correction on RNA-seq differential expression analysis**

(A) Cell-type deconvolution of bulk RNA-seq data from isolated Epcam⁺ cells using MuSiC and the publicly available single-cell RNA-seq dataset GSE124872 as reference. Relative proportions of AT1, AT2, goblet, club, and ciliated epithelial cells are shown for AIR and CS samples. Box-and-whisker plot represents median, interquartile range and minimum-to-maximum values. n=8 mice. Statistical analysis was performed using two-way ANOVA followed by Sidak´s multiple comparisons test, *** $p<0.001$.

(B) MA plot illustrating the effect of correcting for estimated AT2 cell proportions in the differential expression analysis. Each dot represents a gene, displaying the relationship between mean expression and the difference in $\log_2$ fold change between corrected and uncorrected DESeq2 models. The clustering of genes near the x-axis indicates that correction for AT2 proportion had minimal impact on differential gene expression results, consistent with the enrichment of AT2 cells within the Epcam⁺ population.

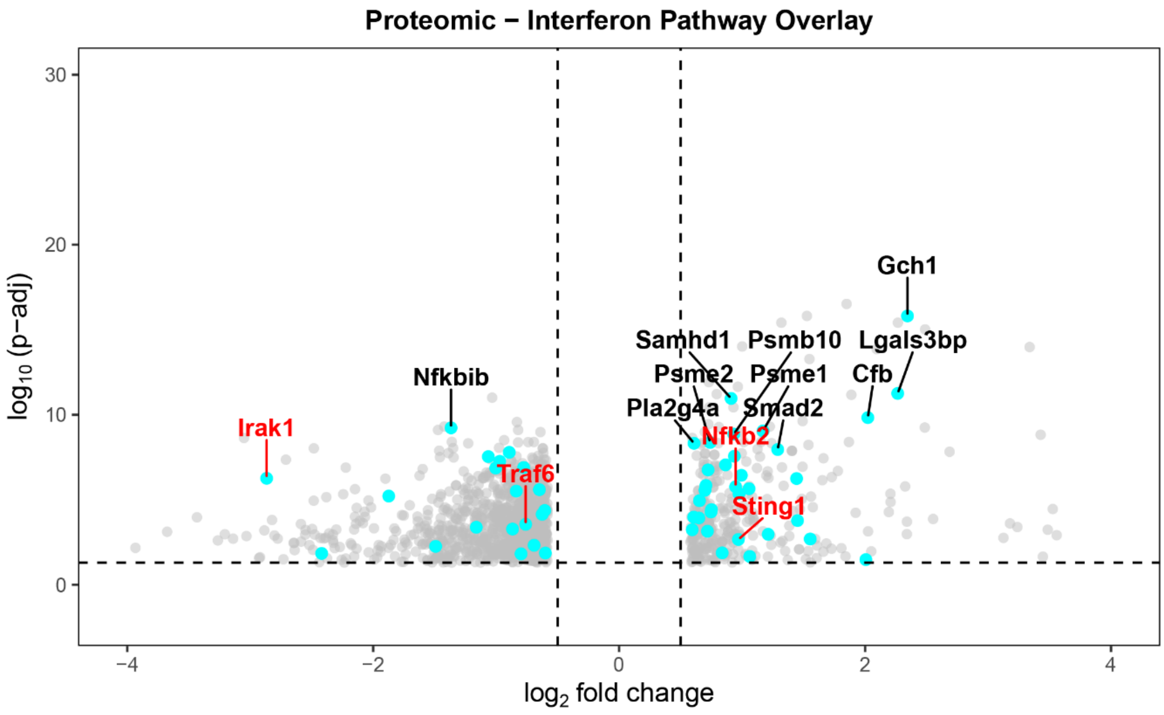


**Supplementary Figure 2: Volcano plot of differentially expressed proteins identified by proteomic analysis.**

Proteomic profiling was performed on Epcam⁺ epithelial cells isolated from lungs of *in vivo* CS-exposed mice, using the same samples as those analyzed by RNA-seq. Proteins were analyzed for differential abundance using thresholds of adjusted p-value (padj < 0.05) and log2 fold change > 0.5. Only significantly regulated proteins were included in the visualization. As only significant proteins are displayed, the central region corresponding to non-significant proteins is not shown. Interferon-related proteins overlapping with the RNA-seq dataset (58 of 380 interferon-associated genes) are highlighted in bold, with key players highlighted in red.

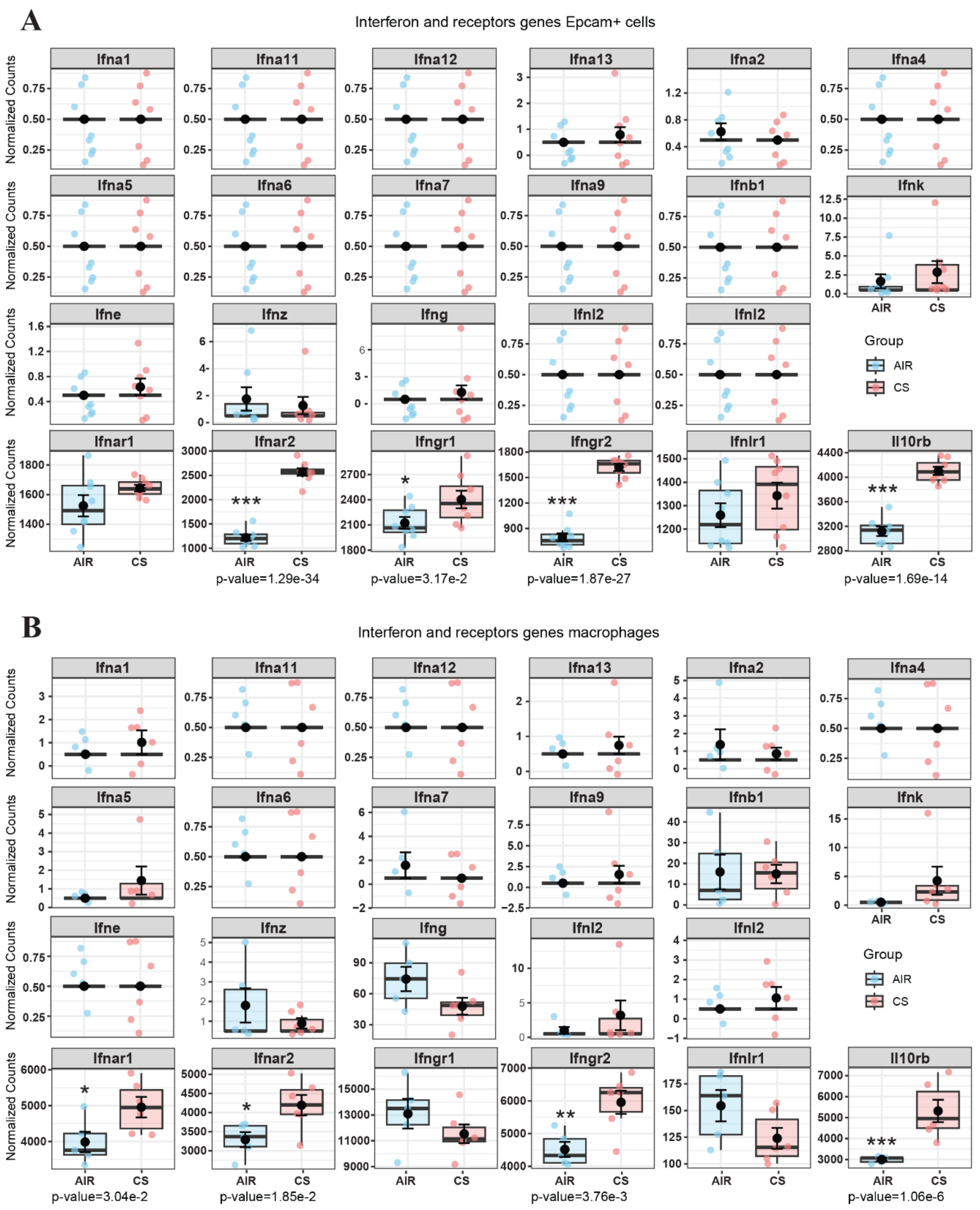


**Supplementary Figure 3: Expression of interferon genes and receptors in epithelial cells and macrophages following cigarette smoke exposure.**

(A) Normalized RNA-seq expression counts of interferon genes and interferon-related receptors in isolated Epcam⁺ epithelial cells from Air and CS-exposed mice. Values represent normalized counts. Data is represented as median with interquartile range and minimum-to-maximum values of n = 8 mice per group. Statistical significance was determined by differential expression analysis (DESeq2), and adjusted p-values are indicated. * $p<0.05$, *** $p<0.001$.

(B) Normalized RNA-seq counts of interferon genes and interferon-related receptors in macrophages isolated from the same mice. Values represent normalized counts. Data is represented as median with interquartile range and minimum-to-maximum values of n = 8 mice per group. Statistical significance was determined by differential expression analysis (DESeq2), and adjusted p-values are indicated. * $p<0.05$, ** $p<0.01$, *** $p<0.001$.

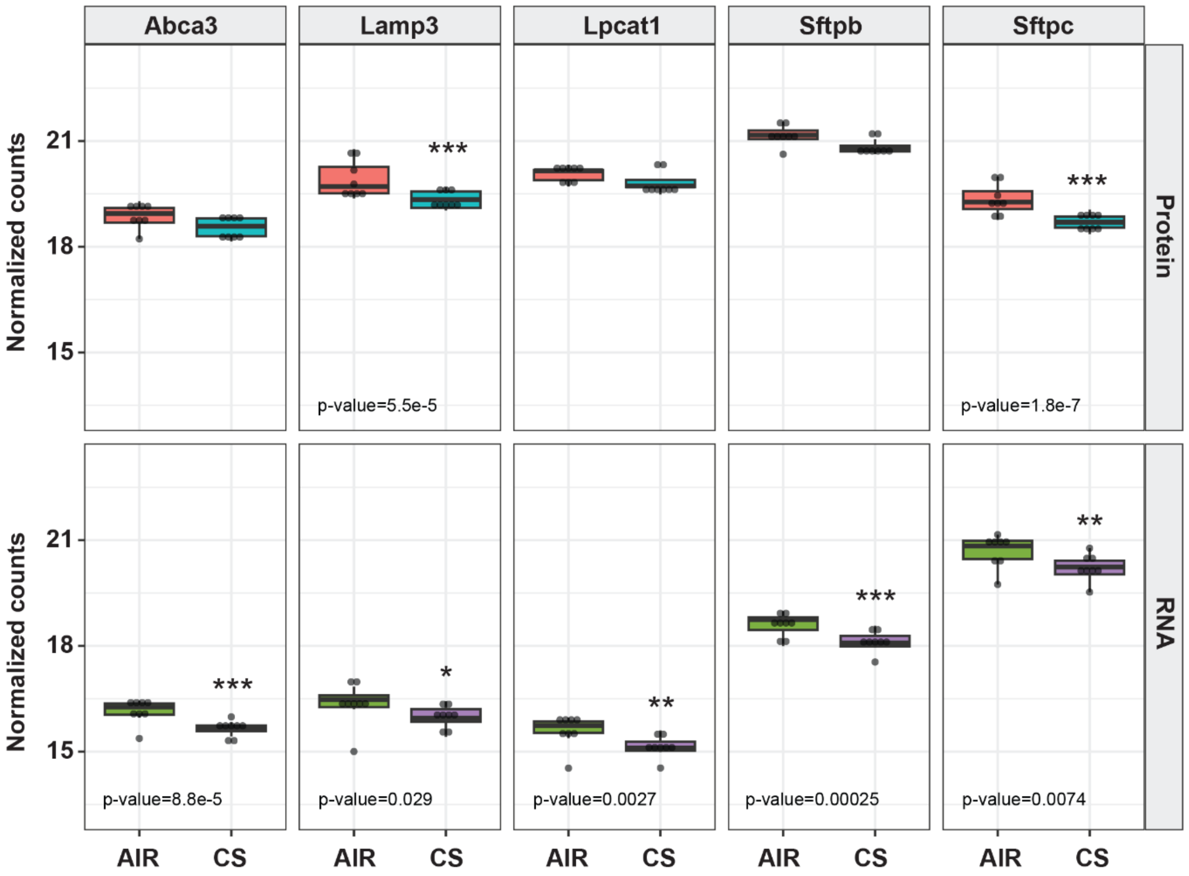


**Supplementary Figure 4: Reduced expression of AT2-associated markers in Epcam⁺ cells following *in vivo* cigarette smoke exposure.**

Normalized expression levels of *Abca3, Lamp3, Lpcat1, Sftpb,* and *Sftpc* measured by RNA-seq and proteomics analyses in isolated Epcam⁺ cells from *in vivo* CS exposure. RNA-seq data are shown as DESeq2-normalized counts, and proteomics data as normalized protein abundance values. Data is represented as median with interquartile range and minimum-to-maximum values of n = 8 mice per group. Statistical significance was determined by differential expression analysis (DESeq2) for RNA-seq data and differential protein expression analysis for proteomics data, and adjusted p-values are indicated. * $p<0.05$, ** $p<0.01$, *** $p<0.001$.

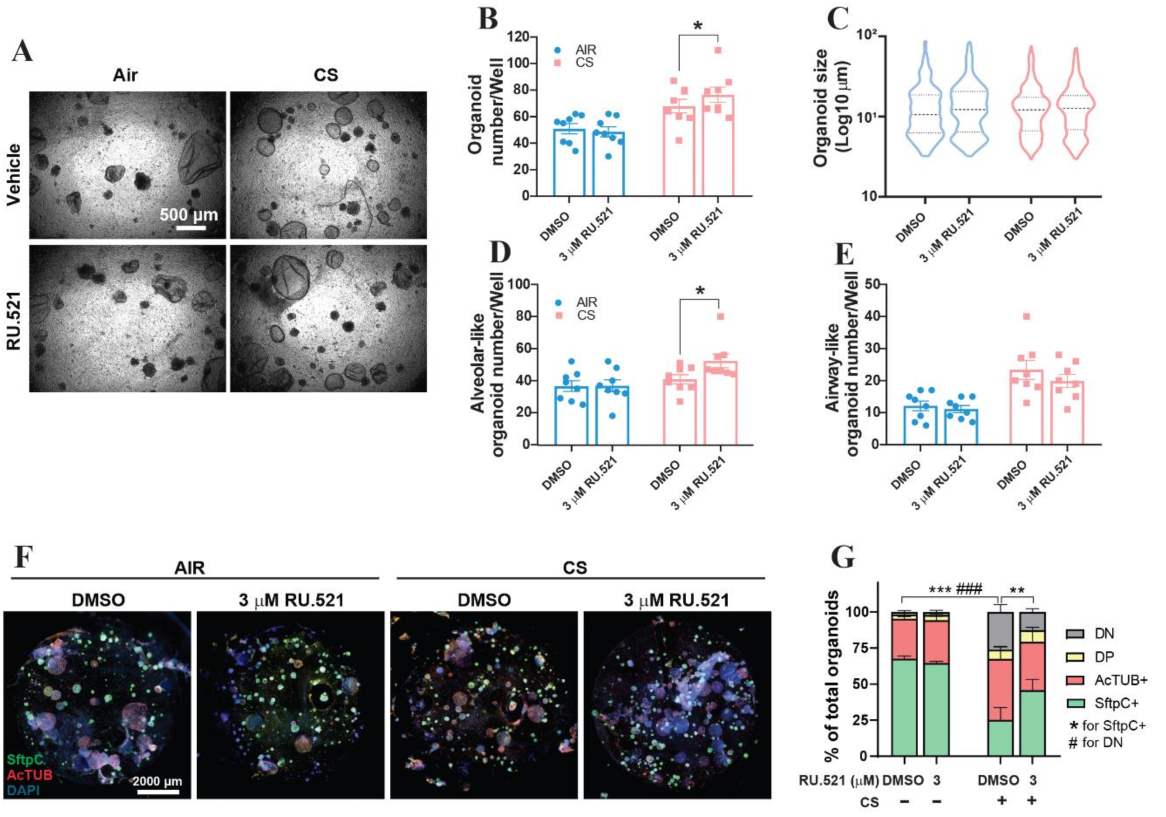


**Supplementary Figure 5: *In vitro* pharmacological inhibition of cGAS selectively enhances alveolar regeneration in cigarette smoke *in vivo*-exposed mice.**

(A) Representative light microscopy images of organoids derived from Air and *in vivo* 5 CS exposed mice subjected to *in vitro* RU.521 cGas inhibitor treatment or vehicle (DMSO) (Scale bar 500 µm).

(B) Quantification of organoid-forming efficiency. Data represent mean ± SEM of n = 8 mice. Statistical analysis was performed using two-way ANOVA followed by Sidak´s multiple comparisons test, *p<0.05.

(C) Quantification of organoid size under the indicated conditions. Data represent median ± quartiles of n = 8 mice. Statistical analysis was performed using Kruskal-Wallis followed by Dunn´s multiple comparisons test.

Quantification of total airway-like (D) and alveolar-like (E) organoids under the indicated conditions. Data represent mean ± SEM of n = 8 mice. Statistical analysis was performed using two-way ANOVA followed by Sidak´s multiple comparisons test, *$p<0.05$.

(F) Representative immunofluorescence images of organoids under the indicated conditions with SftpC$^+$ (green) and AcTub+ (red) markers, counterstained with DAPI (blue). (Scale bar, 2000 µm).

(G) Quantification of percentage of Sftpc$^+$ alveolar organoids, AcTub$^+$ airway organoids, double-positive (DP, SftpC$^+$/AcTub$^+$) and double-negative (DN, SftpC$^-$/AcTub$^-$) organoids. Data represent mean ± SEM of n = 8 mice. Statistical analysis was performed using two-way ANOVA followed by Dunnett´s multiple comparisons test, * indicates differences in SftpC$^+$, # indicates differences in DN. **$p<0.01$, ***/### $p< 0.001$.

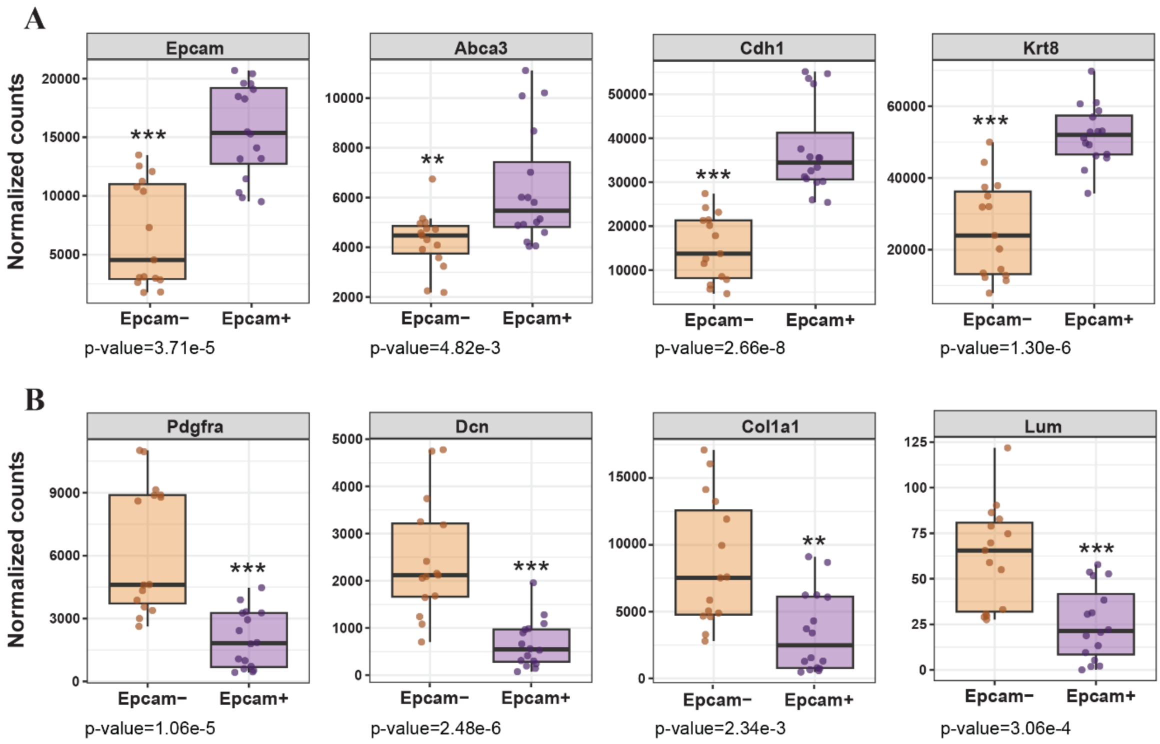


**Supplementary Figure 6: Validation of epithelial and fibroblast enrichment in Epcam⁺ and Epcam⁻ fractions.**

(A) Normalized RNA-seq expression counts of epithelial marker genes in Epcam⁺ and epcam⁻ fractions derived from organoid cultures. Genes include *Epcam, Abca3*, *Cdh1,* and *Krt8.* Values represent normalized counts. Data are shown as individual data points with boxplots representing the median interquartile range and minimum-to-maximum values of n = 16 samples per group. Statistical significance between fractions was assessed using the Wilcoxon rank-sum test on normalized counts per gene, followed by Benjamini–Hochberg correction for multiple testing. Adjusted p-values are indicated. ** p<0.01, *** p<0.001.

(B) Normalized RNA-seq expression counts of fibroblast marker genes in Epcam⁺ and Epcam⁻ fractions derived from the same organoid cultures. Genes include *Pdgfra, Dcn, Col1a1,* and *Lum*. Values represent normalized counts. Data are shown as individual data points with boxplots representing the median and interquartile range and minimum-to-maximum values of n = 16 samples per group. Statistical significance between fractions was assessed using the Wilcoxon rank-sum test on normalized counts per gene, followed by Benjamini–Hochberg correction for multiple testing. Adjusted p-values are indicated. ** $p<0.01$, *** $p<0.001$.

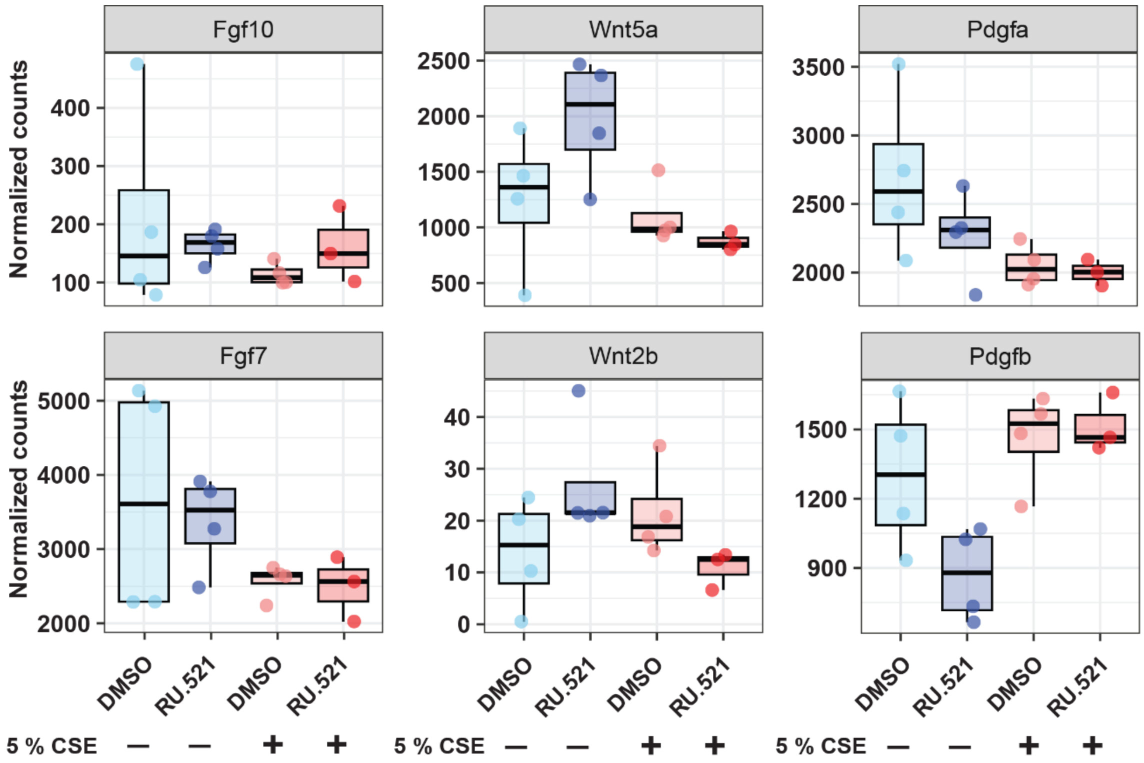


**Supplementary Figure 7: cGAS inhibition does not alter expression of mesenchymal niche factors.**

Normalized expression levels of key mesenchymal support genes (*Fgf10, Fgf7, Wnt5a, Wnt2b, Pdgfa,* and *Pdgfb*) in CD206 murine fibroblast fraction across experimental conditions. Values represent normalized counts. Data is represented as median with interquartile range and minimum-to-maximum values of n = 4 mice per group. Statistical significance was determined by differential expression analysis (DESeq2).
